# Cutaneous inputs from perineal region facilitates and modulates spinal locomotor activity and reduces cutaneous reflexes from the foot in spinal cats

**DOI:** 10.1101/2020.07.29.226530

**Authors:** Angèle N Merlet, Jonathan Harnie, Madalina Macovei, Adam Doelman, Nathaly Gaudreault, Alain Frigon

**Affiliations:** Department of Pharmacology-Physiology, Faculty of Medicine and Health Sciences, Université de Sherbrooke, Sherbrooke, Quebec J1H 5N4, Canada; School of Rehabilitation, Faculty of Medicine and Health Sciences, Université de Sherbrooke, Sherbrooke, Quebec J1H 5N4, Canada; Centre de Recherche du CHUS, Sherbrooke, Quebec J1H 5N4, Canada

**Keywords:** locomotion, sensorimotor interactions, spinal cord injury, spinal reflexes, vibration

## Abstract

It is well known that mechanically stimulating the perineal region potently facilitates hindlimb locomotion and weight support in mammals with a spinal transection (spinal mammals). However, how perineal stimulation mediates this excitatory effect is poorly understood. We evaluated the effect of mechanically stimulating (vibration or pinch) the perineal region on ipsilateral (9-14 ms onset) and contralateral (14-18 ms onset) short-latency cutaneous reflex responses evoked by electrically stimulating the superficial peroneal or distal tibial nerve in seven adult spinal cats where hindlimb movement was restrained. Cutaneous reflexes were evoked before, during, and after mechanical stimulation of the perineal region. We found that vibration or pinch of the perineal region effectively triggered rhythmic activity, unilateral and bilateral to nerve stimulation. When electrically stimulating nerves, adding perineal stimulation modulated rhythmic activity by decreasing cycle and burst durations and by increasing the amplitude of flexors and extensors. Perineal stimulation also disrupted the timing of the ipsilateral rhythm, which had been entrained by nerve stimulation. Mechanically stimulating the perineal region decreased ipsilateral and contralateral short-latency reflex responses evoked by cutaneous inputs, a phenomenon we observed in muscles crossing different joints and located in different limbs. The results suggest that the excitatory effect of perineal stimulation on locomotion and weight support is not mediated by increasing cutaneous reflex gain and instead points to an excitation of central pattern-generating circuitry. Our results are consistent with a state-dependent modulation of reflexes by spinal interneuronal circuits.

**Significance Statement:** Mechanically stimulating the skin of the perineal region strongly facilitates hindlimb locomotion in mammals following a complete spinal cord injury (SCI). Despite its remarkable effectiveness in promoting hindlimb locomotion in spinal cord-injured mammals, we do not know how this is mediated. The present study provides data on how inputs from the perineal region interact with neuronal circuits that generate locomotor-like activity and reflexes from the foot. A better understanding of how inputs from the perineal region interact with neuronal circuits of the spinal cord could lead to non-invasive approaches to restore walking in people with SCI.

## Introduction

Transection of the spinal cord at low thoracic levels completely and permanently abolishes communication between the brain and spinal networks controlling leg movements. Preclinical models of spinal transection (i.e. spinal animals), such as cats, rats and mice, recover hindlimb locomotion with treadmill training (Shurrager & Dykman, 1951; Lovely *et al*., 1986, 1990; Barbeau & Rossignol, 1987; Hodgson *et al*., 1994; Belanger *et al*., 1996; de Leon *et al*., 1998; De Leon *et al*., 1999; Leblond *et al*., 2003; Cha *et al*., 2007; Sławińska *et al*., 2012) or, as shown recently, without task-specific training (Harnie *et al*., 2019). This is due to the presence of a network of neurons located within the lumbar cord that produces the basic hindlimb locomotor pattern, the so-called central pattern generator (CPG) (McCrea & Rybak, 2008; Rossignol & Frigon, 2011; Kiehn, 2016). However, in spinal animals, somatosensory feedback is required to initiate hindlimb locomotion and to adjust the pattern for task demands.

Studies have shown that some types of somatosensory feedback inhibit locomotion while others facilitate it. For example, mechanically stimulating the lumbar region inhibits weight support and locomotion, or locomotor-like activity, in chronic spinal cats in various preparations with the hindlimbs free to move or restrained (Frigon *et al*., 2012; Hurteau *et al*., 2015; Merlet *et al*., 2020) and in curarized decerebrate rabbits with intact or transected spinal cords (Viala & Buser, 1974; Viala *et al*., 1978). In contrast, tonic stimulation of the skin of the perineal region (scrotum, vulva, base of the tail and inguinal fold) facilitates hindlimb locomotion in spinal mammals. For decades, perineal stimulation has been used to initiate fictive locomotion in acute spinal decerebrate curarized preparations treated with clonidine, an α2-noradrenergic agonist (Forssberg & Grillner, 1973; Pearson & Rossignol, 1991; Pearson *et al*., 1992; McCrea *et al*., 1995; Bennett *et al*., 1996) and to facilitate or reinforce hindlimb locomotion in chronic spinal mammals (Barbeau & Rossignol, 1987; Belanger *et al*., 1996; Leblond *et al*., 2003; Langlet *et al*., 2005; Hochman, 2012; Alluin *et al*., 2015; Harnie *et al*., 2019). Stimulating the skin of the perineal region can trigger a well-coordinated pattern of hindlimb locomotion by increasing spinal neuronal excitability through an undefined mechanism (Pearson & Rossignol, 1991). Harnie et al. (2019) proposed that the increase in spinal neuronal excitability provided by tonic perineal stimulation replaces monoaminergic drive from the brainstem. Indeed, several pharmacological studies demonstrated that injecting noradrenergic agonists in spinal cats (Forssberg & Grillner, 1973; Barbeau *et al*., 1987; Chau *et al*., 1998a, 1998b) or serotonergic agonists in spinal mice or rats (Kim *et al*., 2001; Antri *et al*., 2002, 2005; Sławińska *et al*., 2012) facilitates hindlimb locomotion by increasing spinal neuronal excitability. Functionally, the facilitatory effect of perineal region presumably underlies some important survival function, such as escape from a predator (Rossignol *et al*., 2006).

Although we know that activating mechanoreceptors from the perineal region increases the excitability of spinal sensorimotor circuits generating weight support and locomotion, we do not know how this excitatory effect is mediated and how it interacts with the spinal locomotor CPG. One approach to investigate changes in sensorimotor interactions is to evoke and evaluate short-latency cutaneous reflexes from the foot because they project to several motor pools bilaterally via spinal interneurons intercalated between primary afferents and motoneurons (Burke *et al*., 1971; Lundberg *et al*., 1977; Pierrot-Deseilligny *et al*., 1981; Crone *et al*., 1987; Jankowska, 1992). Cutaneous inputs from the foot also exert powerful effects on spinal circuits controlling locomotion (reviewed in Zehr & Stein, 1999; Duysens *et al*., 2000; Rossignol *et al*., 2006).

Therefore, as a first step in determining how perineal stimulation facilitates sensorimotor functions in spinal animals, we investigated how mechanoreceptive inputs from the perineal region interact with short-latency cutaneous reflex responses to ipsilateral and contralateral muscles evoked by electrically stimulating nerves that supply the dorsal and plantar surfaces of the paw in adult chronic spinal cats. We performed two methods of mechanical stimulation of the perineal region: vibration and pinch. We hypothesized that mechanoreceptive inputs from the perineal region mediate their excitatory effect on locomotion and weight support by increasing the gain of short-latency cutaneous reflexes from the foot, thus supplying greater excitatory inputs to spinal sensorimotor circuits.

## Materials and Methods

### Ethical approval

All procedures were approved by the Animal Care Committee of the Université de Sherbrooke in accordance with policies and directives of the Canadian Council on Animal Care (Protocol 442-18). We obtained the current dataset from 7 adult cats (3 females and 4 males) weighing between 3.5 and 4.7 kg. Before and after experiments, cats were housed and fed in a dedicated room within the animal care facility of the Faculty of Medicine and Health Sciences at the Université de Sherbrooke. As part of our effort to maximize the scientific output of each animal, all animals were used in previous studies to answer other scientific questions (Harnie *et al*., 2019; Merlet *et al*., 2020).

### General surgical procedures

We performed implantation and spinal transection surgeries under aseptic conditions in an operating room with sterilized equipment, as described previously (Harnie et al. 2019; Merlet et al. 2020). Briefly, an intramuscular injection containing butorphanol (0.4 mg/kg), acepromazine (0.1 mg/kg), glycopyrrolate (0.01 mg/kg) and ketamine/diazepam (0.11 ml/kg in a 1:1 ratio, i.m.) was performed in cats for sedation and induction before surgery. Cats were then anesthetized with isoflurane (1.5-3%) using a mask and then intubated with a flexible endotracheal tube. Anesthesia was maintained by adjusting isoflurane concentration as needed (1.5-3%). After surgery, we injected an antibiotic (Convenia, 0.1 ml/kg) subcutaneously and taped a transdermal fentanyl patch (25 μg/h) to the back of the animal 2-3 cm rostral to the base of the tail for prolonged analgesia (removed after 5-7 days). We also injected buprenorphine (0.01 mg/kg), a fast-acting analgesic, subcutaneously at the end of the surgery and ~7 h later. After each surgery, we placed the cats in an incubator until they regained consciousness. At the conclusion of the experiments, a lethal dose of pentobarbital was administered through the left or right cephalic vein.

### Electromyography and nerve stimulation

To record the electrical activity of muscles (EMG, electromyography), we directed pairs of Teflon-insulated multistrain fine wires (AS633; Cooner Wire) subcutaneously from two head-mounted 34-pin connectors (Omnetics) that were sewn into the belly of selected hindlimb muscles. To verify electrode placement during surgery, we electrically stimulated each muscle through the appropriate head connector channel to assess the biomechanically desired muscle contraction. The current data set includes EMG from the following muscles: semitendinosus (St, knee flexor/hip extensor), iliopsoas (IP, hip flexor), biceps femoris anterior (BFA, hip extensor), vastus lateralis (VL, knee extensor) and lateral gastrocnemius (LG, ankle extensor/knee flexor). During experiments, EMGs were pre-amplified (10×, custom-made system), band-pass filtered (30-1000 Hz) and amplified (100-5000×) using a 16-channel amplifier (AM Systems Model 3500). EMG data were digitized (5000 Hz) with a National Instruments card (NI 6032E), acquired with custom-made acquisition software, and stored on a computer.

For bipolar nerve stimulation, pairs of Teflon-insulated multistrain fine wires (AS633; Cooner Wire) were passed through a silicon tubing. A horizontal slit was made in the tubing and wires within the tubing were stripped of their insulation. The ends protruding through the cuff were knotted to hold the wires in place and glued. The ends of the wires away from the cuff were inserted into four-pin connectors (Hirose or Samtec) for bipolar nerve stimulation. Cuff electrodes were directed subcutaneously from head-mounted connectors and placed around the left and right superficial peroneal (SP) and tibial (Tib) nerves at ankle level. At this level, the SP nerve is purely cutaneous whereas the Tib nerve is mixed, primarily innervating the plantar skin but also intrinsic foot muscles (Bernard *et al*., 2007).

### Spinal transection and post-lesion period

The spinal cord was completely transected (spinalization) between the 12th and 13^th^ thoracic vertebrae. An incision of the skin was made over the last thoracic vertebrae. After carefully setting aside muscle and connective tissue, a small laminectomy of the dorsal bone was made to expose the spinal cord. Lidocaine (xylocaine) was applied topically and two to three injections were made within the spinal cord, which was then transected with surgical scissors. Hemostatic material (Spongostan) was inserted within the gap and muscles and skin were sewn back to close the opening in anatomic layers. After spinalization, the bladder was manually expressed one to two times daily and cats were monitored by experienced personnel. To confirm that the spinal transection was complete in all cats, we performed histological analysis and qualitative and quantitative evaluations of the lesioned area, as described and shown in Harnie et al. (2019).

As previously described (Merlet *et al*., 2020), cats of the present study were used by Harnie *et al*. (2019) to determine the effects of different interventions on the recovery of hindlimb locomotion after spinal transection. One week after the spinal transection, cats were divided into three groups. Group 1 (Cats 3 and 4), received manual therapy five times a week for 5 weeks, 20 min per day. Briefly, cats were placed on a padded support and a cushioned restraint stabilized the pelvis and an experimenter applied rhythmic unilateral pressure to the triceps surae muscles on each leg for 10 min. Group 2 (Cats 5, 7 and 8) received locomotor training on a treadmill five times a week for 5 weeks, 20 min per day. For locomotor training, two experimenters manually moved the hindlimbs to simulate locomotion with appropriate joint kinematics and paw contacts on a motorized treadmill at a speed of 0.4 m/s with the forelimbs placed on a stationary platform. Group 3 (Cats 10 and 11) did not receive any treatment for the 6 weeks after spinal transection. The experiments of the present study were performed in 6 of those cats (Cats 3, 4, 5, 7, and 10, 11; 3 females and 3 males) with perineal vibration and 7 cats (Cats 3, 4, 5, 7, 8, 10, 11; 3 females and 4 males) with perineal pinch.

### Experimental protocol

**Figure 1** schematically illustrates the experimental set-up and protocol. As previously described (Merlet *et al*., 2020), we positioned the cat on a cushion and a padded support restrained the pelvic region from movement. We secured one hindpaw to a stationary restraint while the other hindpaw was secured to a robotic arm using medical tape. The robotic arm was stationary and used only to position the limb during experiments. Hip, knee and ankle joints were positioned at ~120°, 90° and 90°, respectively, for both hindlimbs (**Fig. 1A**).

**Figure 1.**
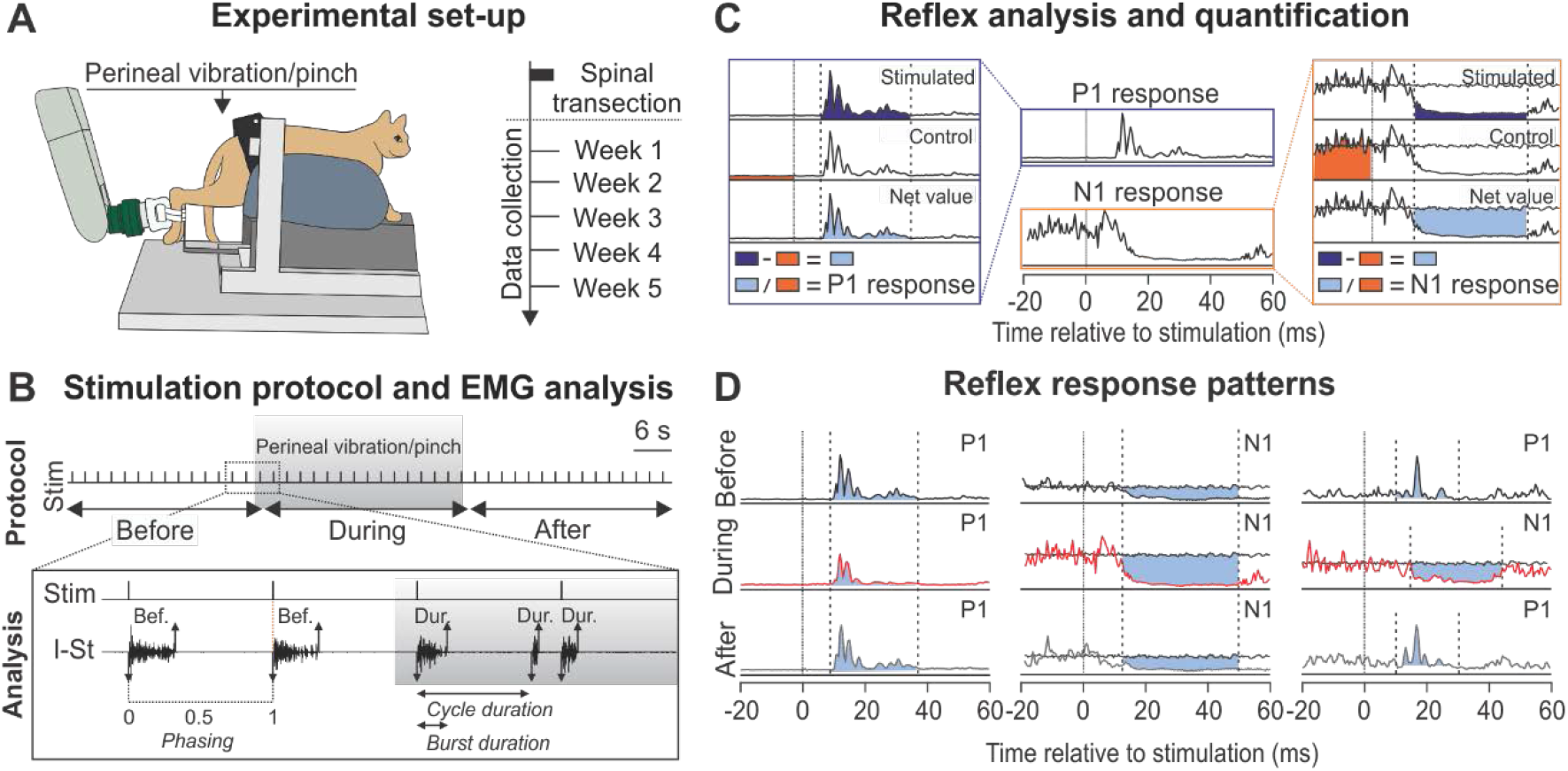
Experimental design. ***A***: experimental setup for reflex testing. Data were collected each week after spinal transection for 5 weeks (W). ***B*** (*top*): during a trial, the superficial peroneal and the tibial nerve were stimulated for 90 s at 0.5 Hz (1 every 2 sec), for a total of ~45 stimuli. Vibration or pinch of the perineal region was initiated 30 after the beginning of the trial for 30 s. ***B*** (*bottom*): we measured some electromyography (EMG) parameters, such as cycle and burst durations as well as EMG phasing with the nerve stimulation (Stim), before, during and after the vibration/pinch. We determined burst onsets (downward arrows) and offsets (upward arrows) in selected studied muscles. ***C***: stimulated cycles were sorted into periods (before, during and after the vibration/pinch) and time windows were determined to quantify short-latency excitatory (P1) and inhibitory (N1) responses. ***D***: time windows in some reflex responses patterns. The onset and offset of the time windows were the same within a trial, except when we observed a switch from a P1 to an N1 response, where we adjusted time windows (*right*).

During experiments, we evoked cutaneous reflexes by delivering electrical stimuli to the SP or Tib nerves with a Grass S88 Stimulator at an intensity of 1.2 times the motor threshold, defined as the voltage required to elicit a small consistent short-latency (8-10 ms) excitatory response in an ipsilateral flexor, such as St. Each stimulation consisted of a train of three 0.2 ms pulses delivered at 300 Hz. In each condition, we delivered ~45 stimuli for 90 s at 0.5 Hz (1 train every 2 s), i.e. ~15 stimulations were given for each period (**Fig. 1B**, *top*). The vibration was delivered using a custom device with a 120-Hz frequency and an amplitude of 1.5 mm applied over the perineal region. The most effective site to perform perineal stimulation was the skin of the scrotum in males and around the vulva in females (Sherrington, 1910). The pinch of the perineal skin was performed by rubbing it between the thumb, index, and middle finger at a constant moderate to strong pressure. The same experimenter (J. Harnie) presented all stimuli to all cats. We collected all data (i.e. SP and Tib nerve stimulation and vibration and pinch of the perineal region) within a single session for a given cat and each week after the spinal transection for 5 weeks.

### Data analysis

#### EMG analysis and quantification

We measured some EMG parameters in each period of the trial in selected muscles. Only trials where rhythmic bursts of the selected muscles observed before, during and after the perineal stimulation were selected (*n* = 9-14 trials). To quantify EMG bursts, we selected a flexor (IP) and an extensor (LG) muscle. An experimenter (Merlet) determined burst onsets and offsets of selected muscles by visual inspection from the raw EMG waveforms using a custom-made program. Cycle duration was measured as the time between two successive bursts and burst duration was determined as the time from onset to offset (**Fig. 1B**, *bottom*). We characterized mean EMG amplitude by integrating the full-wave EMG burst from onset to offset and dividing it by its burst duration. We also measured the interval of time between flexor burst onset and the first electrical stimulus of the train normalized to cycle duration, termed the phasing (**Fig. 1B**, *bottom*). We also measured the phasing between ipsilateral and contralateral flexor burst onsets.

#### Reflex analysis and quantification

The step-by-step procedure for quantifying reflex responses is illustrated in **Fig. 1C**. Reflex responses were sorted into 3 periods: before, during and after the vibration/pinch of the perineal region and averaged (*n* = 13-15 nerve stimulations per period). For each nerve stimulation, we averaged the 20 ms window preceding the electrical stimulation to provide a baseline EMG (blEMG) in each period. The blEMG provides an indication of the level of excitability of the motor pool when the afferent volley arrives from the SP or Tib nerve electrical stimulation. To measure reflex amplitude, we set a window from the onset to the offset of the response, defined as prominent negative or positive deflections away from the blEMG. We used previous studies in intact and spinal cats as a reference for setting the windows (Duysens & Loeb, 1980; Abraham *et al*., 1985; Loeb, 1993; Frigon & Rossignol, 2008; Hurteau *et al*., 2017, 2018; Hurteau & Frigon, 2018). Based on the terminology introduced by Duysens and Loeb (1980), we defined short-latency (~8-10 ms) excitatory (P1) or inhibitory responses (N1) and mid-latency (~20-25 ms) excitatory responses (P2). We also classified responses in contralateral muscles as P1 or N1 because they had onsets <18 ms, which is the minimal latency for reflexes with a relay in the brainstem in cats (Shimamura & Livingston, 1963). In spinal cats, the longer-latency response (P2) starting at ~20-25 ms is small or absent (Frigon & Rossignol, 2008; Hurteau *et al*., 2017; Hurteau & Frigon, 2018; Merlet *et al*., 2020). In the present study, we observed some P2 responses, in particular in IP and LG muscles, but not enough to perform statistical analyses. Therefore, we analyzed ipsilateral and contralateral short-latency responses with onset latencies of ~9-14 ms and ⁓14-18 ms, respectively.

The window for the short-latency excitatory (P1) or inhibitory (N1) responses started when the averaged EMG was greater than the blEMG for ≥ 3 ms using 95% confidence intervals and with a minimal latency of 7 ms. Onset latencies had to be adjusted because response onset differed slightly depending on the muscle and between cats. The window ended when the reflex response returned to blEMG for ≥ 3 ms or at a maximal latency of ~40 ms for some muscles. The onset and offset of the windows were the same for each period of the trial. When we observed the appearance of N1 or P1 responses during perineal stimulation with no response observed before or after, we placed a window with identical onsets and offsets for comparison. When we observed a switch from a P1 response to an N1 response within a trial, we adjusted the onset and the offset of the window at each period (**Fig. 1D**).

The EMG of P1 or N1 responses was integrated and the blEMG was subtracted to provide a net reflex value. This net value was then divided by the control value to evaluate reflex responses independently of the level of EMG activity (**Fig. 1C**) (Matthews, 1986; Frigon & Rossignol, 2007, 2008; Hurteau *et al*., 2017, 2018; Hurteau & Frigon, 2018; Merlet *et al*., 2020). The division is necessary to determine whether reflex amplitude scales with blEMG. For example, if the stimulated EMG had a value of 2 arbitrary units (a.u.) with a blEMG of 4 a.u. before the stimulation, this would give a net reflex value of −2 a.u. (2 a.u. - 4 a.u.) and a normalized value of −0.5 a.u. (−2 a.u./4 a.u.) with the division. If blEMG doubled with the perineal vibration or pinch to 8 a.u. and the stimulated EMG also doubled to 4 a.u., then the net reflex value would be −4 a.u. (4 a.u. - 8 a.u.), or twice the value of 2 a.u. obtained without the application of the perineal vibration or pinch. However, with the division, the normalized value is equal to −0.5 a.u. (−4 a.u./8 a.u.), indicating that the reflex response simply scaled with blEMG. We defined a response as absent (i.e. no response) when the net reflex value was between −0.1 a.u. and 0.1 a.u. To determine whether P1 or N1 responses were modulated with mechanical stimulation of the perineal region, responses in a given muscle were expressed as a percentage of the before period value. As we observed excitatory responses (P1) during a period and inhibitory responses (N1) during another period within a trial, the modulation of reflex responses with the mechanical stimulation of the perineal region sometimes exceeded 100%.

#### Video recordings

During experiments, one camera (Basler AcA640-100 gm) captured video at 60 frames/s with a spatial resolution of 640 by 480 pixels. A custom-made Labview program acquired images and synchronized them with the EMG to determine the initiation and termination of mechanical stimulation of the perineal region with vibration or pinch. We used custom-made software to analyze videos offline at 60 frames/s.

### Statistics

To evaluate the effects of mechanically stimulating the perineal region with vibration and pinch on EMG bursts and on electrically-evoked cutaneous reflexes, we performed a one-factor [period (before, during, after)] repeated-measures ANOVA. When a main effect was found, a *post hoc* analysis was conducted using Fisher’s test. To compare the effects of mechanical stimulation of the perineal region on EMG bursts bilaterally, we performed paired *t* tests (ipsilateral *vs* contralateral). The normality of each variable was assessed by the Shapiro Wilk test. As previously described for reflex analysis (Merlet *et al*., 2020), we pooled data across trials collected over 5 weeks after spinal transection because we found no visible effect of week after spinal transection on the amplitude of reflex responses. To demonstrate this, we measured the modulation of reflexes evoked by SP or Tib stimulation during vibration or pinch of the perineal region each week after spinal transection in the ipsilateral St. We found no significant effect of week on the modulation of reflexes (*p* > 0.05, repeated measures ANOVA; data not shown), suggesting that the modulation in reflexes with perineal stimulation was not affected as a function of time after spinal transection. Thus, each trial was treated separately and data were pooled for statistical comparisons, with a single cat contributing up to 5 trials. Statistical analyses on reflex responses were performed with a minimum of 9 trials in at least 2 cats. The critical level for statistical significance was set at an α-level of 0.05. Analyses were done with Statistica 8 (Statsoft, Tulsa, OK, USA). All values in the test and figures are the mean ± standard deviation (SD).

## Results

### Modulation of locomotor-like activity with mechanical stimulation of the perineal region

In chronic spinal cats with their hindlimbs restrained, electrical stimulation of the SP or Tib nerves generates rhythmic bursts of activity in multiple hindlimb muscles (Merlet *et al*., 2020). To determine how perineal stimulation affected these bursts, we recorded hindlimb EMG before, during and after mechanically stimulating the perineal region, as shown in **Figure 2**. Note that the EMGs for a given muscle are at the same vertical scale within a trial. On the right of each condition, we present the average rectified EMG trace before, during and after the perineal stimulation of ipsilateral selected muscles over the cycle normalized to a hindlimb flexor ipsilateral to the nerve stimulation. In all the trials shown, we observed alternating bursts of activity between ipsilateral flexors (I-St and I-IP) and extensors (I-LG and I-VL) before the mechanical stimulation of the perineal region. In most cases, SP or Tib nerve stimulation initiated bursts in ipsilateral flexors, whereas ipsilateral extensor activity occurred spontaneously but ceased with the following nerve stimulation that initiated another flexor burst. In other words, electrical nerve stimulation reset the extension phase to flexion and entrained the rhythm. On the other hand, contralateral rhythmic activity was weaker and less frequent with nerve stimulation alone. With the addition of perineal stimulation, we observed more frequent rhythmic alternating activity in flexor and extensor muscles unilateral to the nerve stimulation, with greater amplitude, which was not necessarily timed to the nerve stimulation. Perineal stimulation consistently increased burst amplitude, with the exception of the I-St (**Fig. 2**, *rightmost panels*). Upon cessation of perineal stimulation, the activity of flexor and extensor muscles returned to that observed before perineal stimulation.

**Figure 2.**
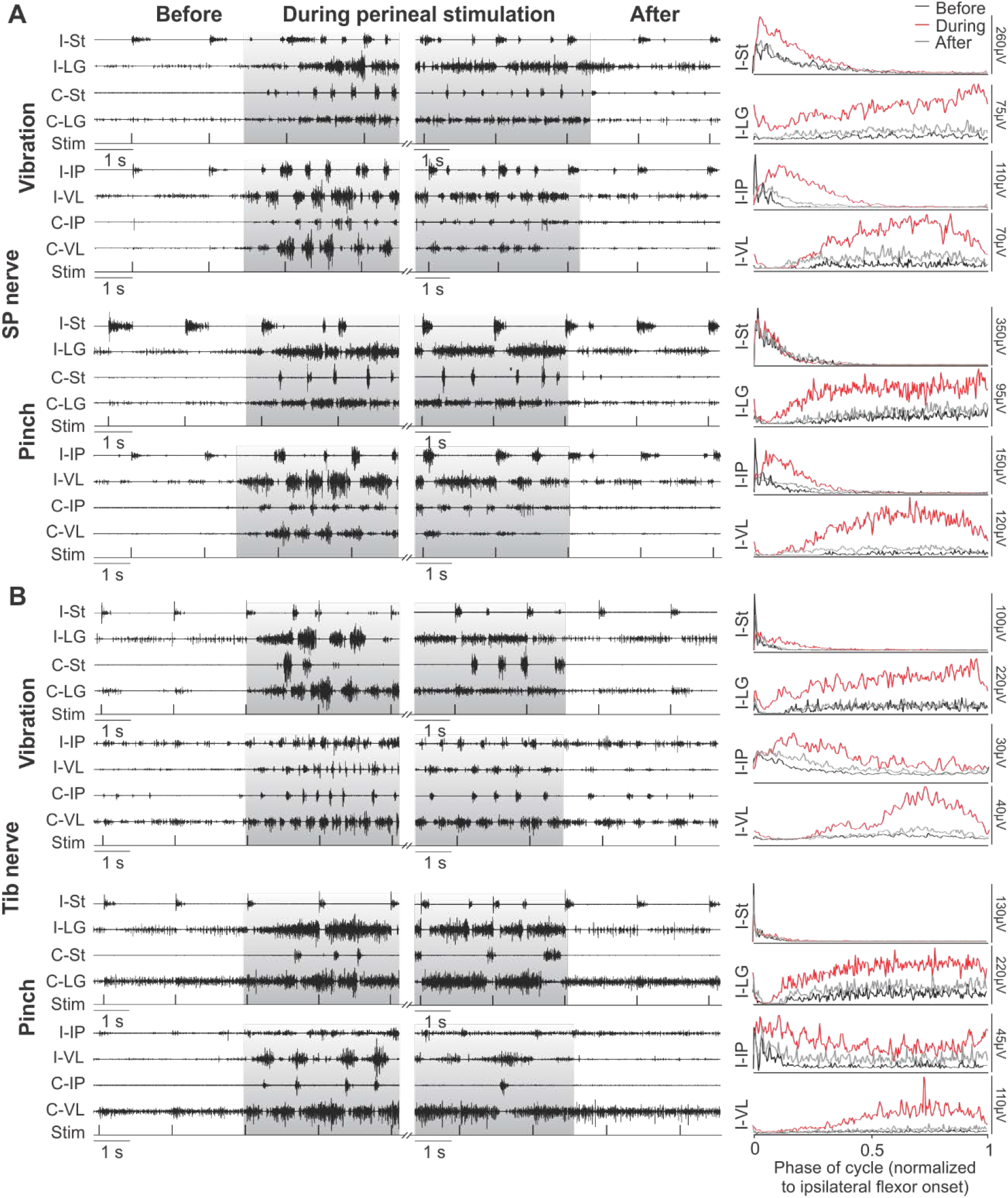
Modulation of hindlimb muscle activity with mechanical stimulation of the perineal region. The left of each panel shows the electromyography (EMG) from the ipsilateral (I) and contralateral (C) semitendinosus (St), lateralis gastrocnemius (LG), iliopsoas (IP) and vastus lateralis (VL) while stimulating the superficial peroneal (SP) (***A***) or tibial (Tib) (***B***) nerves before, during (in grey) and after the vibration or pinch of the perineal region. The timing of the stimulation (Stim) is shown below the EMGs. The right of each panel shows the average rectified EMG traces of selected ipsilateral muscles normalized to burst onset before, during and after perineal stimulation. Data are from Cat 4 at week 2 after spinal transection.

**Table 1** shows the presence of rhythmic activity unilateral and bilateral to SP and Tib nerve stimulation before, during, and after vibration or pinch of the perineal region for pooled data (i.e. the sum of trials over 5 weeks). Note that bilateral activity means an out-of-phase alternation between the ipsilateral and contralateral sides for a given muscle. Before perineal stimulation, we observed alternating bursts of activity between flexors and extensors unilaterally (29-40% of the trials) and bilaterally (11-33% of the trials). Perineal stimulation increased the presence of rhythmic activity unilaterally (70-83% of trials) and bilaterally (67-74% of trials). When vibration/pinch of the perineal region was removed, the presence of rhythmic activity was slightly elevated compared to the before period (unilateral: 43-49% and bilateral; 17-33%). It is important to note that perineal stimulation rarely induced rhythmic activity in Cats 7 and 8. Interestingly, these cats did not recover forward hindlimb locomotion after spinal transection (Harnie *et al*., 2019).

**Table 1.**
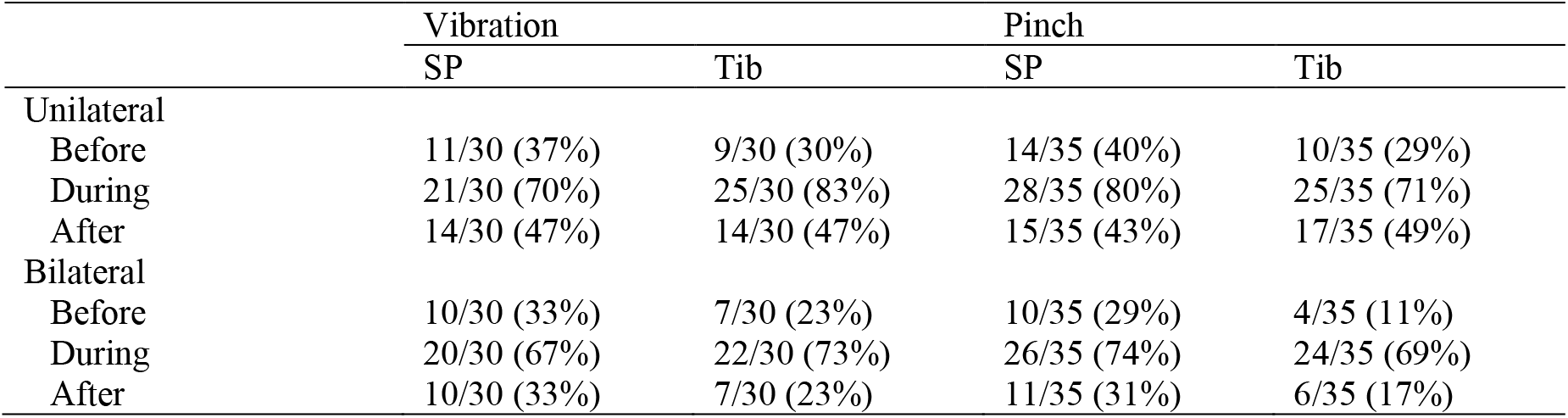
Presence of rhythmic activity patterns with stimulation of SP and Tib nerves before, during and after vibration or pinch of the perineal region. Values are the sum of trials over 5 weeks when we observed locomotor-like activity unilateral and bilateral to the nerve stimulations represented as a fraction and a percentage (in parentheses) of the total number of trials. SP, superficial peroneal nerve; Tib, tibial nerve.

|  | Vibration |  | Pinch |  |
| --- | --- | --- | --- | --- |
|  | SP | Tib | SP | Tib |
| Unilateral |  |  |  |  |
| Before | 11/30 (37%) | 9/30 (30%) | 14/35 (40%) | 10/35 (29%) |
| During | 21/30 (70%) | 25/30 (83%) | 28/35 (80%) | 25/35 (71%) |
| After | 14/30 (47%) | 14/30 (47%) | 15/35 (43%) | 17/35 (49%) |
| Bilateral |  |  |  |  |
| Before | 10/30 (33%) | 7/30 (23%) | 10/35 (29%) | 4/35 (11%) |
| During | 20/30 (67%) | 22/30 (73%) | 26/35 (74%) | 24/35 (69%) |
| After | 10/30 (33%) | 7/30 (23%) | 11/35 (31%) | 6/35 (17%) |

To quantify the effects of mechanically stimulating the perineal region on rhythmic activity, we measured cycle and burst durations as well as mean EMG amplitude in an ipsilateral flexor (I-IP) and extensor (I-LG) muscle before, during and after perineal stimulation. For these analyses, we selected only trials where we observed rhythmic activity before, during and after perineal stimulation (*n* = 9-14 trials). We assessed EMG changes using a one-way repeated measures ANOVA. In I-IP and I-LG muscles, vibration of the perineal region significantly decreased the cycle duration in trials with SP (F_(2,20)_ = 5.41, *p* = 0.013, −20%; F_(2,18)_ = 5.51, *p* = 0.014, −26%, respectively) and Tib nerve (F_(2,24)_ = 14.50, *p* = 7.4×10^−5^, −25%; F_(2,22)_ = 14.91, *p* = 8.1×10^−5^, −31%, respectively) stimulations (**Fig. 3A**, *left panels*). Pinch of the perineal region also significantly decreased the cycle duration of I-IP and I-LG muscles while stimulating the Tib nerve (F_(2,26)_ = 14.87, *p* = 5.0×10^−5^, −25%; F_(2,16)_ = 4.98, *p* = 0.021, −15%, respectively) but not for the SP nerve (F_(2,26)_ = 0.08, *p* = 0.927; F_(2,24)_ = 1.94, *p* = 0.166, respectively) (**Fig. 3A**, *right panels*). Perineal vibration significantly decreased the burst duration of I-IP and I-LG muscles with SP nerve stimulation (F_(2,20)_ = 7.08, *p* = 0.005, −18%; F_(2,18)_ = 4.25, *p* = 0.031, −16%, respectively) and of I-LG (F_(2,24)_ = 1.16, *p* = 0.330; F_(2,22)_ = 4.67, *p* = 0.020, −30%, respectively) with Tib nerve stimulation (**Fig. 3B**, *left panels*). Perineal pinch only decreased the burst duration of I-IP while stimulating the Tib nerve (F_(2,26)_ = 3.94, *p* = 0.032, −25%; F_(2,16)_ = 3.33, *p* = 0.062, respectively) whereas with SP nerve stimulation, burst durations of I-IP and I-LG muscles were unchanged (F_(2,26)_ = 2.46, *p* = 0.105; F_(2,24)_ = 1.87, *p* = 0.175, respectively) (**Fig. 3B**, *right panels*). Perineal vibration significantly increased the mean EMG amplitude of I-IP and I-LG muscles while stimulating the SP (F_(2,20)_ = 13.54, *p* = 1.9×10^−4^, +85%; F_(2,18)_ = 17.93, *p* = 5.2×10^−5^, +171%, respectively) and Tib nerves (F_(2,24)_ = 12.02, *p* = 2.4×10^−4^, +93%; F_(2,22)_ = 19.26, *p* = 1.5×10^−5^, +209%, respectively) (**Fig. 3C**, *left panels*). Perineal pinch also significantly increased the mean EMG amplitude of I-IP and I-LG muscles while stimulating the SP (F_(2,26)_ = 23.05, *p* = 1.7×10^−6^, + 112%; F_(2,24)_ = 39.20, *p* = 2.7×10^−8^, +180%, respectively) and Tib nerves (F_(2,26)_ = 64.76, *p* = 8.0×10^−11^, +212%; F_(2,16)_ = 30.58, *p* = 3.4×10^−6^, +462%, respectively) (**Fig. 3C**, *right panels*). Thus, in general, vibration and pinch of the perineal region increased the frequency of the rhythm by reducing the duration of flexor and extensor bursts while increasing their amplitudes.

**Figure 3.**
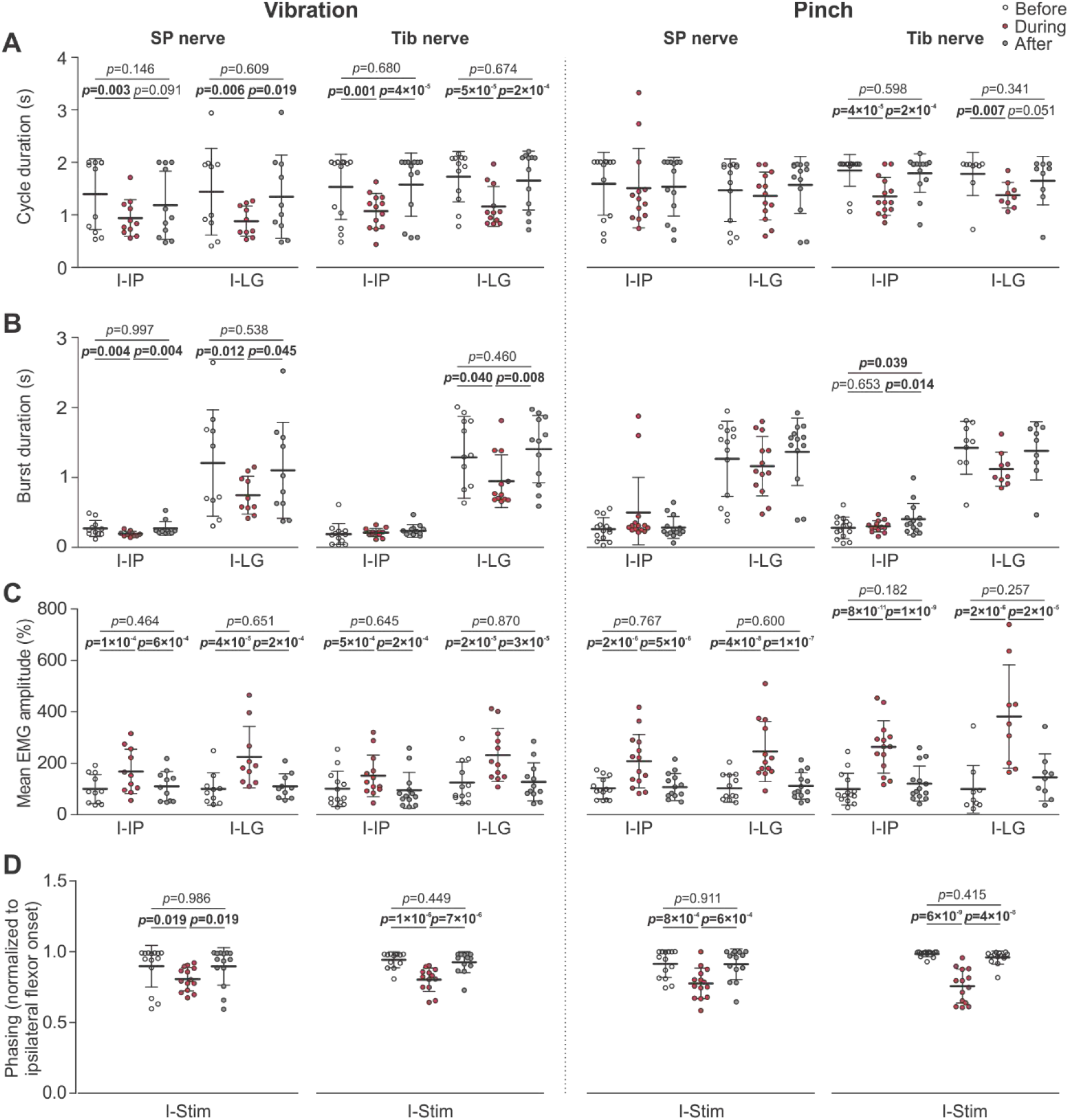
Modulation of EMG parameters in ipsilateral hindlimb muscles with mechanical stimulation of the perineal region. Each panel shows the cycle (***A***) and burst (***B***) duration as well as the mean electromyography (EMG) amplitude (***C***) in the ipsilateral iliopsoas (I-IP) and lateralis gastrocnemius (I-LG) muscles while stimulating the superficial peroneal (SP) or tibial (Tib) nerves before, during and after vibration or pinch of the perineal region. **D)** Phasing of I-IP between the flexor burst onset and the first electrical stimulus of the train. Scatterplots show the mean ± SD from pooled data (*IP* and *LG*: SP-vibration *n* = 11 and 10 trials from 4 and 4 cats, respectively; SP-pinch *n* = 14 and 13 trials from 3 and 4 cats, respectively; Tib-vibration *n* = 13 and 12 trials from 3 and 4 cats, respectively; Tib-pinch *n* =14 and 9 trials from 3 and 3 cats, respectively). *p* values comparing periods were obtained with Fisher’s post-hoc test if a significant main effect of the ANOVA was found.

To quantify if ipsilateral flexor bursts remained timed to the nerve stimulation during perineal stimulation, we measured the phasing of I-IP to SP and Tib nerve stimulations before and after perineal stimulation. Before perineal stimulation, the phasing was ~1, indicating that nerve stimulation initiated the flexor burst. We found that the phasing of I-IP was significantly reduced during vibration (F_(2,20)_ = 4.96, *p* = 0.027, −7%; F_(2,24)_ = 34,98, *p* = 1.4×10^−6^, −15%, respectively) and pinch (F_(2,26)_ = 12.38, *p* = 7.0×10^−4^, −14%; F_(2,26)_ = 43.41, *p* = 5.4×10^−9^, −23%, respectively) of the perineal region, with a value of ~0.8 (**Fig. 3D**), indicating that flexor burst onset was less timed to or entrained by the nerve stimulation. After perineal stimulation, the phasing returned to ~1.

To evaluate the effects of mechanically stimulating the perineal region on bilateral rhythmic activity, we compared cycle and burst durations of ipsilateral and contralateral IP and LG muscles as well as the phasing of I-IP and C-IP muscles during perineal stimulation. The rhythms ipsilateral and contralateral to the nerve stimulation showed different types of relationships, with 1 or more cycles occurring on one side relative to the other, as occurs during split-belt locomotion on the ‘slow’ and ‘fast’ sides (Forssberg *et al*., 1980; Frigon *et al*., 2017). Thus, we also report 1:1, 1:2, 1:3, 1:4, 1:5 and 1:6 relationships in percentage of the total number of IP cycles, indicating that the fast limb had one, two, three, four, five and six cycles for every cycle of the slow limb (**Fig. 4**). For these analyses, we selected only trials where out-of-phase activity between the ipsilateral and contralateral sides was observed during perineal stimulation (*n* = 14-18 trials). In the four EMGs trials shown, we observed out-of-phase activity between I-IP and C-IP (**Fig. 4A**, *top*) with a phasing of ~0.4 (**Fig. 4A**, *bottom left*). Note that a phasing of 0.5 represents a strict out-of-phase left-right alternation. In some cases, we observed different patterns of rhythmic activity between I-IP and C-IP. A relationship of 1:3 is shown during pinch of the perineal region with Tib nerve stimulation indicating that the fast limb (I-IP) had three cycles for every cycle of the slow limb (C-IP). In this case, we observed that the last cycle of the fast limb (I-IP) was the longest cycle and that the duration and amplitude of extensor burst (I-BFA, *biceps femoris anterior*) was increased. A relationship of 1:1 predominated in all conditions but during pinch of the perineal region with Tib nerve stimulation, we reported a substantial proportion of other relationships (1:2, 1:3, 1:4, 1:5 and 1:6) (~40%) compared to the other conditions (~20%) (**Fig. 4A**, *bottom right*). Note that 1:2+ relationships were observed in 76-88% of the trials. We reported that cycle duration was significantly longer in the contralateral LG compared to the ipsilateral LG during pinch of the perineal region while stimulating the Tib nerve (t_(15)_ = −2.37, *p* = 0.033, +40%) whereas no significant differences were found for the other conditions (SP-vibration t_(16)_ = −0.76, *p* = 0.460; SP-pinch t_(17)_ = 1.00, *p* = 0.333; Tib-vibration t_(14)_ = −1.46, *p* = 0.168) or for the IP muscle (SP-vibration t_(14)_ = 1.62, *p* = 0.129; SP-pinch t_(17)_ = −1.85, *p* = 0.082; Tib-vibration t_(17)_ = −1.62, *p* = 0.124; Tib-pinch t_(14)_ = −2.12, *p* = 0.054) (**Fig. 4B**). The burst duration of the contralateral IP was significantly greater compared to the ipsilateral IP while stimulating the Tib nerve during vibration (t_(17)_ = −2.23, *p* = 0.040, +25%) and pinch (t_(14)_ = −2.25, *p* = 0.042, +111%) whereas no significant differences were found while stimulating the SP nerve (t_(14)_ = −1.82, *p* = 0.092; t_(17)_ = 0.30, *p* = 0.764, respectively) (**Fig. 4C**). The burst duration of the contralateral LG was also longer compared to the ipsilateral LG during perineal pinch while stimulating the Tib nerve (t_(15)_ = −2.15, *p* = 0.049, +49%) whereas no significant differences was reported in the other conditions (SP-vibration t_(16)_ = −0.94, *p* = 0.360; SP-pinch t_(17)_ = 0.88, *p* = 0.390; Tib-vibration t_(14)_ = −1.62, *p* = 0.129) (**Fig. 4C**).

**Figure 4.**
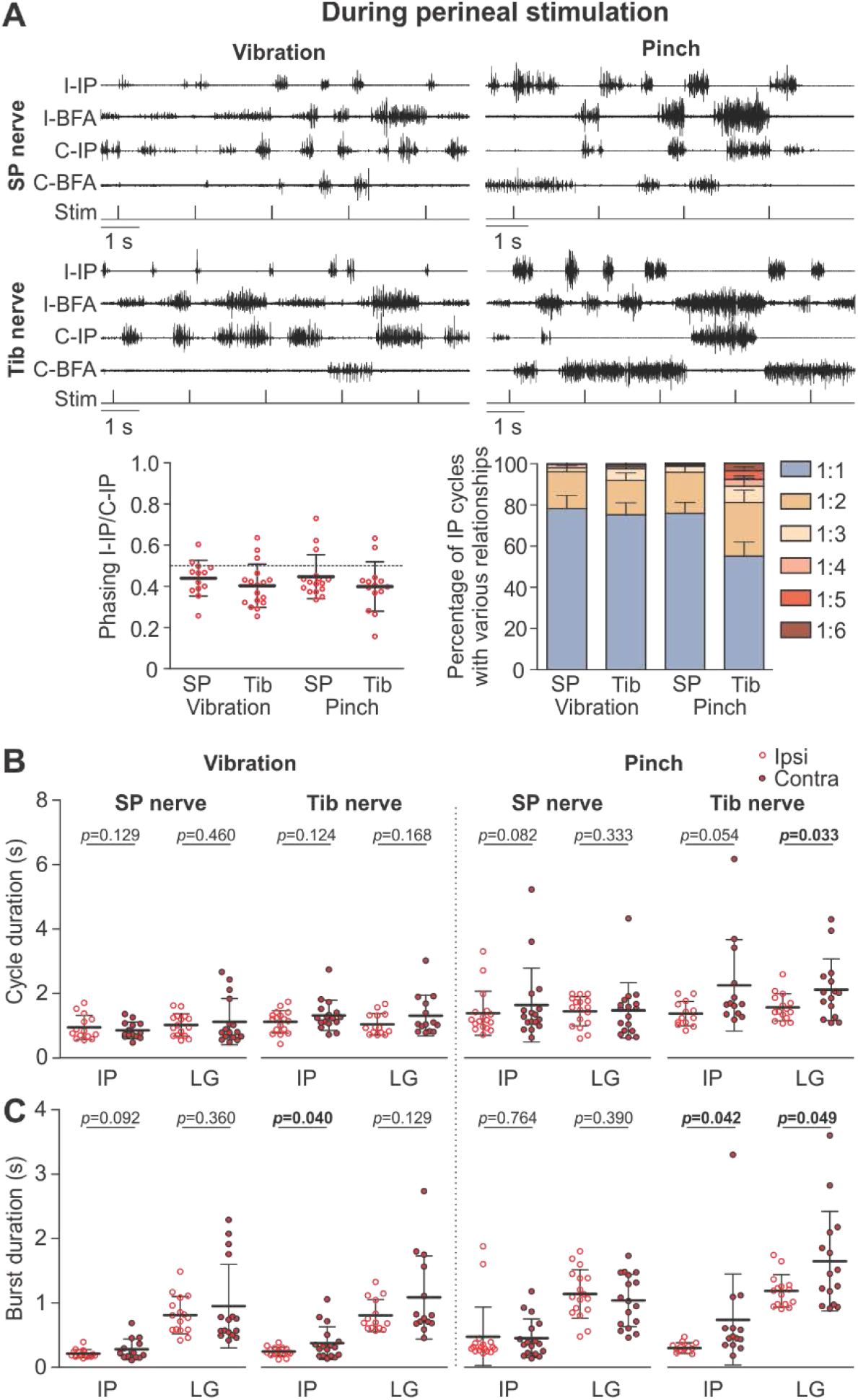
Modulation of hindlimb muscle activity during mechanical stimulation of the perineal region. ***A***: The top panel shows electromyography from the ipsilateral (I) and contralateral (C) iliopsoas (IP) and biceps femoris anterior (BFA) with stimulation (Stim) of the superficial peroneal (SP) or tibial (Tib) nerves during vibration or pinch of the perineal region. Data are from Cat 11 at week 4 after spinal transection. The panel below shows the phasing between I-IP and C-IP (*left*) and the percentage of IP cycles with 1:1, 1:2, 1:3, 1:4, 1:5 and 1:6 relationships (*right*) in all conditions. Each panel shows the cycle (***B***) and burst (***C***) durations of the ipsilateral and contralateral IP and lateralis gastrocnemius (LG) muscles during vibration/pinch of the perineal region. Bar graphs or scatterplots show the mean ± SD from pooled data (*IP* and *LG*: SP-vibration *n* = 14 and 16 trials from 4 and 5 cats, respectively; SP-pinch *n* = 17 and 17 trials from 4 and 4 cats, respectively; Tib-vibration *n* = 18 and 14 trials from 4 and 4 cats, respectively; Tib-pinch *n* = 14 and 15 trials from 3 and 4 cats, respectively). *p* values comparing ipsilateral and contralateral sides were obtained with paired *t* tests.

In summary, tonic stimulation of the perineal region with vibration or pinch is effective in facilitating or triggering unilateral (~76% of the trials) and bilateral (~71% of the trials) rhythmic activity in spinal cats. Adding perineal stimulation to SP and Tib nerve stimulations modulates the existing rhythmic activity by decreasing cycle and burst durations and increasing mean EMG amplitude in ipsilateral flexors and extensors. Perineal stimulation also produced different patterns of rhythmic activity between the ipsilateral and contralateral sides, as the sensory feedback from the tonic perineal stimulation competes with phasic inputs from nerve stimulations to elicit rhythmic activity.

### Modulation of cutaneous reflexes with mechanical stimulation of the perineal region

We investigated the effects of mechanically stimulating the perineal region on cutaneous reflexes from the paws in spinal cats. We electrically stimulated the SP and Tib nerves before, during, and after applying vibration or pinch of the perineal region and recorded reflex responses in ipsilateral and contralateral hindlimb muscles. We set the stimulation intensity at 1.2 times the motor threshold to elicit a small consistent response in an ipsilateral flexor (St), which has the lowest activation threshold. However, this stimulation intensity did not result in reflex responses in all recorded muscles. **Table 2** summarizes the presence of responses in the four selected hindlimb muscles.

**Table 2.**
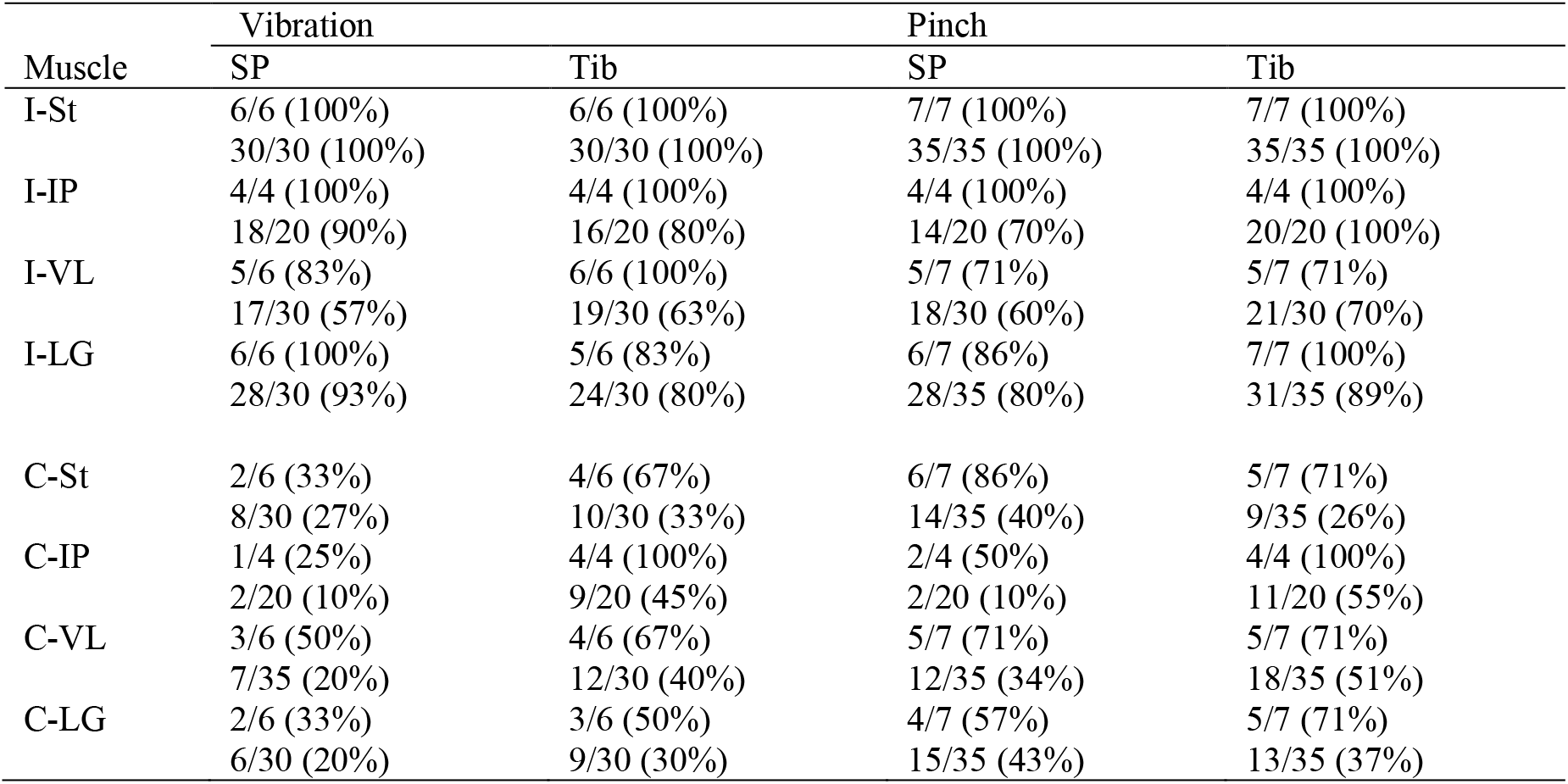
Presence of reflex responses evoked by the stimulation of SP or Tib nerves during vibration or pinch of the perineal region. For each muscle, the number of cats where reflex responses were observed is represented as a percentage of the total number of cats available. Below, the sum of trials where reflex responses were observed is represented as a percentage of the total number of trials in all cats. SP, superficial peroneal nerve; Tib, tibial nerve.

To facilitate comparisons in reflex responses between conditions and muscles, we organized the figures in the same way (**Figs. 5–8**). Each figure represents a different muscle and panels A and B (or C and D for contralateral responses) show reflex responses evoked by SP and Tib nerve stimulation, respectively. On the left of each subpanel, we showed average traces before, during and after vibration or pinch of the perineal region in an 80-ms window, 20 ms before and 60 ms after the nerve stimulation. Within a trial, we observed short-latency excitatory (P1) or inhibitory (N1) responses, no response (−) or a combination of these responses. The ‘no response’ indicates a value between −0.1 a.u. and 0.1 a.u., even though a small deflection might be observed. For each condition, we show the different types of observed reflex response patterns. Note that reflex responses for a given pattern is on the same vertical scale before, during and after perineal stimulation. To show how N1 responses deviated from the baseline EMG (i.e. the EMG before the stimulation), we superimposed averaged traces that received a stimulation (line in black, red or grey) with averaged traces without stimulation (line in dark grey). In the middle of each panel, we report the occurrence of the reflex response pattern as a percent, representing the percentage of time we observed this pattern within a condition. The amplitude of reflex responses before, during and after the vibration/pinch of the perineal region is shown on the right of each panel for pooled data. As we observed excitatory (P1) and/or inhibitory (N1) responses within a trial, the modulation of reflex responses could exceed 100% in some muscles. In the following paragraphs, the *p* values reported are those of the main effect from a one-way repeated measure ANOVA.

**Figure 5.**
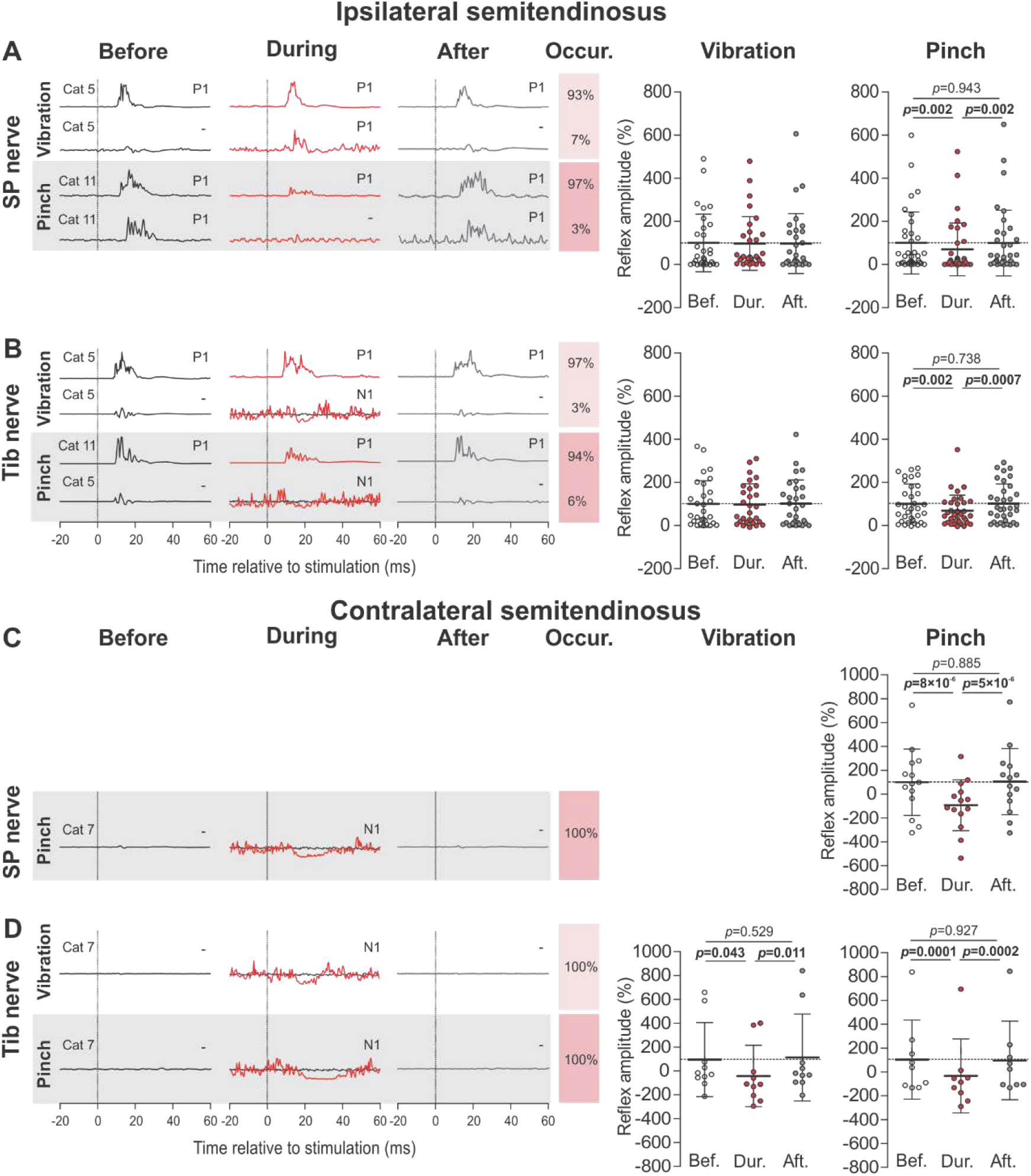
Modulation of reflex responses in the ipsilateral and contralateral semitendinosus muscle by mechanical stimulation of the perineal region. Short-latency ipsilateral and contralateral reflex responses evoked by stimulating the superficial peroneal (SP) (panels ***A*** and ***C***) and tibial (Tib) nerves (***B*** and ***D***, respectively). *Left*: All reflex responses patterns are illustrated for each condition. For each pattern, waveforms are averages of 13-15 stimulations per period for a representative cat in an 80-ms window. To better visualize N1 responses, we superimposed averaged traces that received a stimulation (line in black, red or grey) with averaged traces without stimulation (line in dark grey). On the right of each pattern, the occurrence (occur.) of the reflex responses pattern in percentage are illustrated. *Right*: the effect of vibration and pinch of the perineal region are illustrated for each period (Bef., before; Dur., during; Aft., after). Scatterplots show the mean ± SD of short-latency responses from pooled data (ipsilateral SP-vibration *n* = 30 trials from 6 cats; ipsilateral and contralateral SP-pinch *n* = 35 and 14 trials from 7 and 6 cats, respectively; ipsilateral and contralateral Tib-vibration *n* = 30 and 10 trials from 6 and 4 cats, respectively; ipsilateral and contralateral Tib-pinch *n* = 35 and 10 trials from 7 and 5 cats, respectively). *p* values comparing periods were obtained with Fisher’s post-hoc test if a significant main effect of the ANOVA was found.

We first evaluated the modulation of ipsilateral and contralateral reflex responses with perineal stimulation in the St muscle, a knee flexor and hip extensor (**Fig. 5**). In the ipsilateral St, we predominantly observed short-latency excitatory responses (P1) evoked by SP and Tib nerve stimulation before, during and after the perineal stimulation in 93-97% of the trials. In 3-7% of the other trials, we reported three other types of patterns (**Fig. 5A-B**, *left*). Pinch of the perineal region significantly decreased reflex responses evoked by SP (F_(2,68)_ = 7.00, *p* = 0.002, −30%; **Fig. 5A**, *right*) and Tib nerve (F_(2,68)_ = 7.64, *p* = 0.001, −33%; **Fig. 5B**, *right*) stimulations, whereas vibration did not significantly modulate reflex responses evoked by SP and Tib nerve stimulation (F_(2,58)_ = 0.08, *p* = 0.809; F_(2,58)_ = 0.42, *p* = 0.657, respectively; **Fig. 5A-B**, *right*).

In the contralateral St, stimulating the SP nerve with vibration did not evoke enough reflex responses to perform statistical analysis. In the other conditions, perineal stimulation caused the appearance of N1 responses with SP or Tib nerve stimulation in 100% of the trials when no response (−) was observed before and after (**Fig. 5C-D**, *left*). Thus, in these cases, the appearance of an N1 response results in a reduction of reflex responses with our measure. Pinch of the perineal region significantly decreased reflex responses in the contralateral St evoked by SP nerve stimulation (F_(2,26)_ = 21.14, *p* = 3.5×10^−6^, −193%; **Fig. 5C**, *right*). Vibration and pinch of the perineal region significantly decreased reflex responses evoked by Tib nerve stimulation (F_(2,18)_ = 4.35, *p* = 0.029, −129%; F_(2,16)_ = 16.41, *p* = 1.3×10^−4^, −131%, respectively; **Fig. 5D**, *right*).

We next evaluated the modulation of ipsilateral and contralateral reflex responses by perineal stimulation in the IP muscle, a hip flexor (**Fig. 6**). In the ipsilateral IP, stimulating the SP and Tib nerves predominantly evoked P1 responses before, during and after perineal pinch in 71-100% of the trials, whereas in the other trials, N1 responses could replace the P1 response during the perineal stimulation or the N1 could be present in all three periods (**Fig. 6A-B**, *right*). Pinch of the perineal region significantly decreased reflex responses evoked by SP (F_(2,26)_ = 8.46, *p* = 0.001, −53%; **Fig. 6A**, *right*) and Tib nerve (F_(2,38)_ = 7.17, *p* = 0.002, −55%; **Fig. 6B**, *right*) stimulation whereas vibration did not significantly modulate reflex responses evoked by SP and Tib nerve stimulation (F_(2,34)_ = 1.99, *p* = 0.193; F_(2,30)_ = 2.24, *p* = 0.124, respectively; **Fig. 6A-B**, *right*).

**Figure 6.**
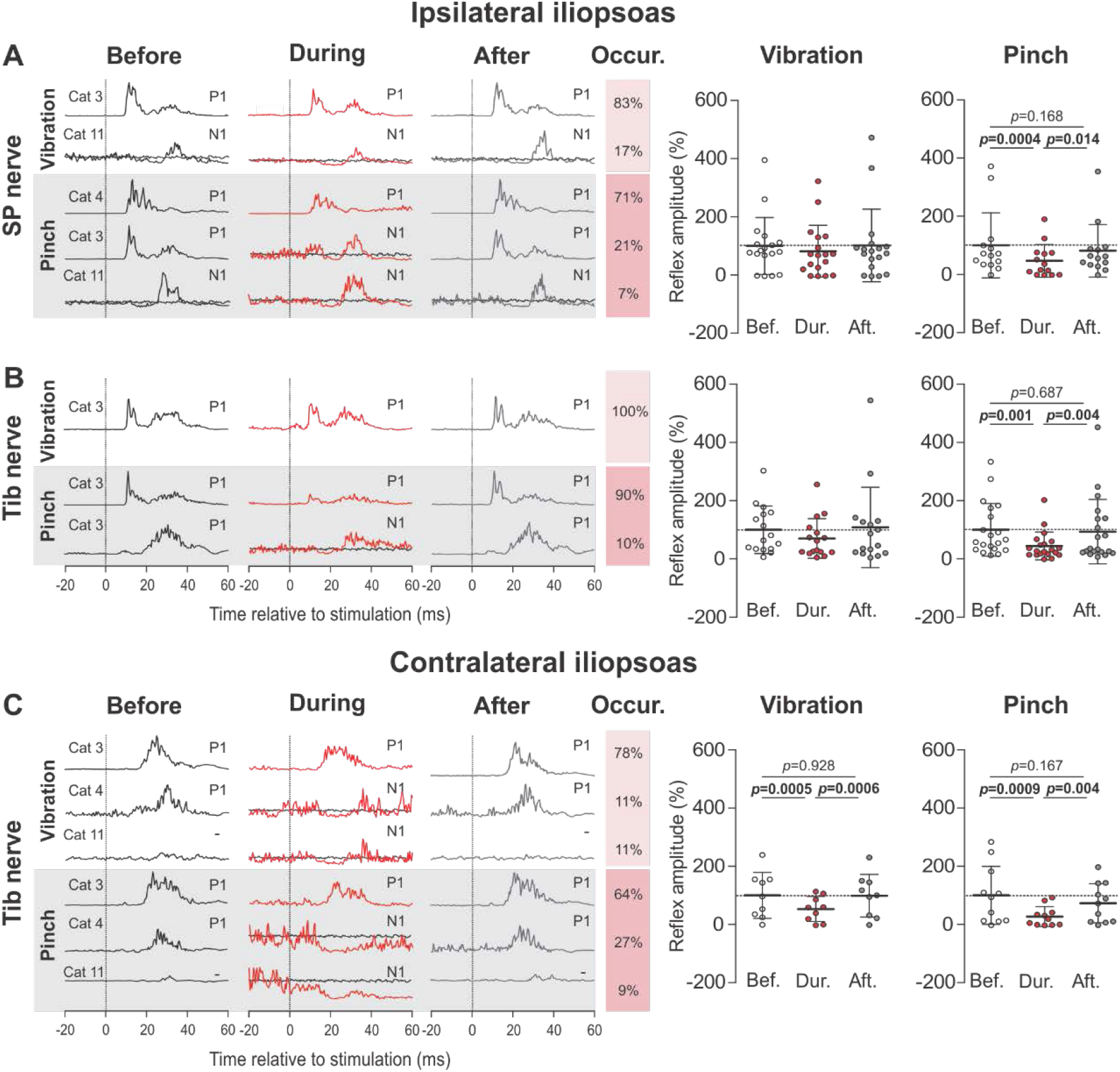
Modulation of reflex responses in the ipsilateral and contralateral iliopsoas muscle by mechanical stimulation of the perineal region. Short-latency ipsilateral and contralateral reflex responses evoked by stimulating the superficial peroneal (SP) (panels ***A*** and ***C***) and tibial (Tib) nerves (***B*** and ***D***, respectively) were showed. *Left*: All reflex responses patterns are illustrated for each condition. For each pattern, waveforms are averages of 13-15 stimulations per period for a representative cat in an 80-ms window. To better visualize N1 responses, we superimposed averaged traces that received a stimulation (line in black, red or grey) with averaged traces without stimulation (line in dark grey). On the right of each pattern, the occurrence (occur.) of the reflex responses pattern in percentage are illustrated. *Right*: the effect of vibration and pinch of the perineal region are illustrated for each period (Bef., before; Dur., during; Aft., after). Scatterplots show the mean ± SD of short-latency responses from pooled data (ipsilateral SP-vibration *n* = 18 trials from 4 cats; ipsilateral SP-pinch *n* = 14 trials from 4 cats, respectively; ipsilateral and contralateral Tib-vibration *n* = 16 and 9 trials from 4 and 4 cats, respectively; ipsilateral and contralateral Tib-pinch *n* = 20 and 11 trials from 4 and 4 cats, respectively). *p* values comparing periods were obtained with Fisher’s post-hoc test if a significant main effect of the ANOVA was found.

In the contralateral IP, we observed reflex responses evoked by Tib nerve stimulation only. Stimulating the Tib nerve mainly evoked contralateral P1 responses before, during and after perineal stimulation (64-78% of the trials). In the other trials, perineal stimulation induced a switch from a P1 response to an N1 response (11-27% of the trials) or an N1 response appeared during perineal stimulation when no response was observed before or after (9-11% of the trials) (**Fig. 6C**, *left*). Vibration and pinch of the perineal region significantly decreased reflex responses evoked by Tib nerve stimulation (F_(2,16)_ = 12.30, *p* = 0.001, −47%; F_(2,20)_ = 7.80, *p* = 0.003, −73%, respectively) (**Fig. 6C**, *right*).

We next evaluated the modulation of ipsilateral and contralateral reflex responses by perineal stimulation the VL muscle, a knee extensor (**Fig. 7**). In the ipsilateral VL, we observed P1 (6-47% of trials) or N1 (11-28% of trials) responses evoked by SP and Tib nerve in the three periods, as well as the appearance of N1 responses (12-61% of trials) or disappearance of P1 responses (11-33%) during perineal stimulation (**Fig. 7A-B**, left). Pinch of the perineal region significantly decreased reflex responses evoked by SP nerve stimulation (F_(2,34)_ = 9.10, *p* = 0.001, −93%) whereas vibration did not significantly affect reflex responses (F_(2,32)_ = 1.44, *p* = 0.253; **Fig. 7A**, *right*). Vibration and pinch of the perineal region significantly decreased reflex responses evoked by Tib nerve stimulation (F_(2,36)_ = 9.05, *p* = 0.007, −94%; F_(2,40)_ = 12.72, *p* = 0.005, −100%, respectively; **Fig. 7B**, *right*).

**Figure 7.**
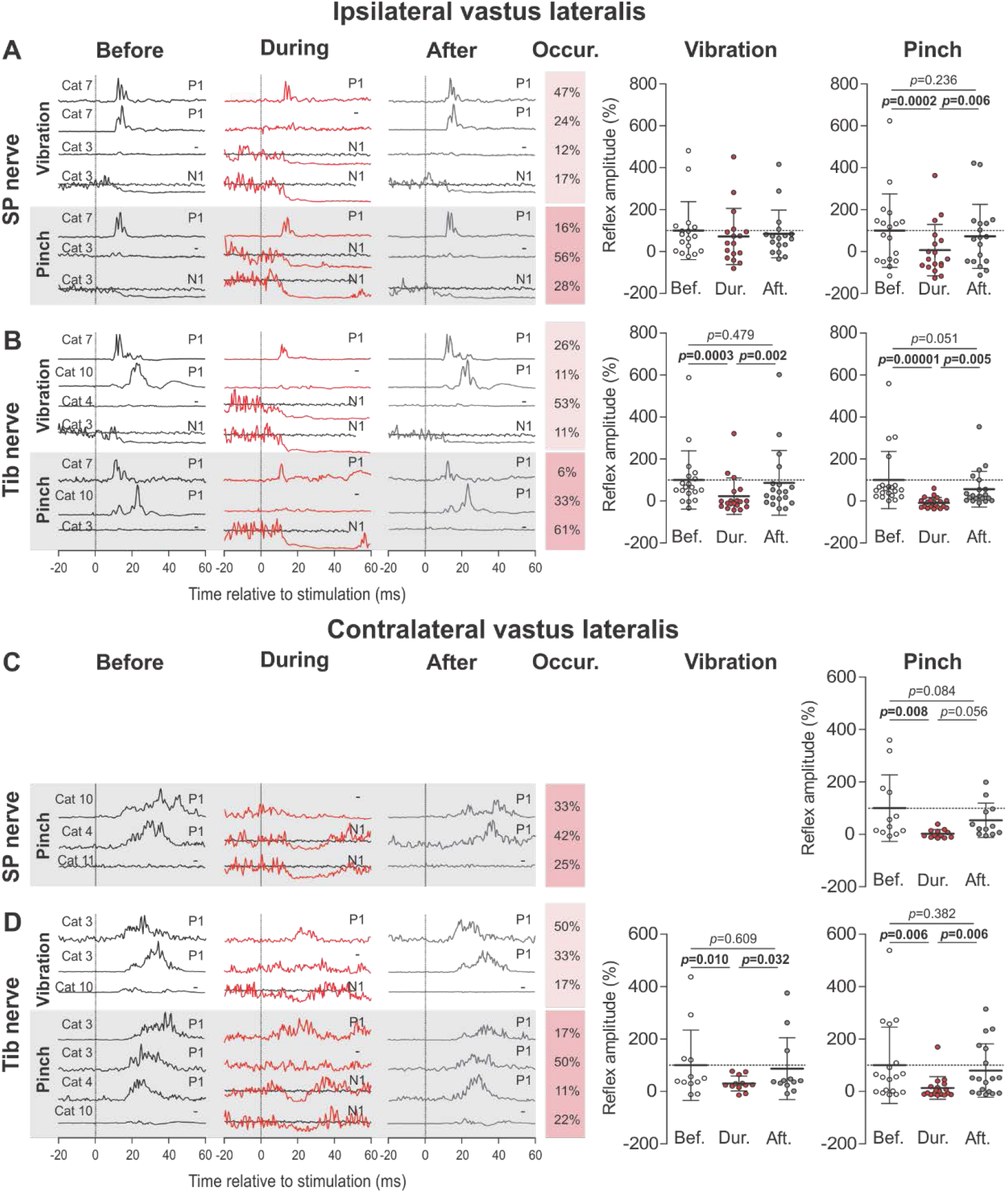
Modulation of reflex responses in the ipsilateral and contralateral vastus lateralis muscle by mechanical stimulation of the perineal region. Short-latency ipsilateral and contralateral reflex responses evoked by stimulating the superficial peroneal (SP) (panels ***A*** and ***C***) and tibial (Tib) nerves (***B*** and ***D***, respectively) were showed. *Left*: All reflex responses patterns are illustrated for each condition. For each pattern, waveforms are averages of 13-15 stimulations per period for a representative cat in an 80-ms window. To better visualize N1 responses, we superimposed averaged traces that received a stimulation (line in black, red or grey) with averaged traces without stimulation (line in dark grey). On the right of each pattern, the occurrence (occur.) of the reflex responses pattern in percentage are illustrated. *Right*: the effect of vibration and pinch of the perineal region are illustrated for each period (Bef., before; Dur., during; Aft., after). Scatterplots show the mean ± SD of short-latency responses from pooled data (ipsilateral SP-vibration *n* = 17 trials from 5 cats; ipsilateral and contralateral SP-pinch *n* = 18 and 13 trials from 5 and 5 cats, respectively; ipsilateral and contralateral Tib-vibration *n* = 19 and 12 trials from 6 and 4 cats, respectively; ipsilateral and contralateral Tib-pinch *n* = 21 and 18 trials from 5 and 5 cats, respectively). *p* values comparing periods were obtained with Fisher’s post-hoc test if a significant main effect of the ANOVA was found.

In the contralateral VL, stimulating the SP nerve during vibration did not evoke enough reflex responses to perform statistical analysis. In the other conditions, we observed P1 responses (17-50% of trials) in the three periods, a switch from P1 to N1 responses during perineal pinch (11-42% of trials), as well as the appearance of N1 (17-25% of trials) or disappearance of P1 responses (33-50% of trials) during perineal stimulation (**Fig. 7C-D**, left). Pinch of the perineal region significantly decreased reflex responses evoked by SP nerve stimulation (F_(2,22)_ = 7.59, *p* = 0.003, −98%; **Fig. 7C**, *right*). Vibration and pinch of the perineal region significantly decreased reflex responses evoked by Tib nerve stimulation (F_(2,22)_ = 4.48, *p* = 0.023, −70%; F_(2,34)_ = 7.91, *p* = 0.002, −87%, respectively; **Fig. 7D**, *right*).

Lastly, we evaluated the modulation of ipsilateral and contralateral reflex responses by perineal stimulation in the LG muscle, an ankle extensor and a knee flexor (**Fig. 8**). In the ipsilateral LG, we observed P1 (23-82% of trials) or N1 (7-25%) responses evoked by SP and Tib nerve stimulation in all three periods or a switch from P1 to N1 responses (11-54% of trials) (**Fig. 8A-B**, *left*). Vibration and pinch of the perineal region significantly decreased reflex responses evoked by SP (F_(2,54)_ = 5.58, *p* = 0.002, −48%; F_(2,54)_ = 9.70, *p* = 2.5×10^−4^, − 75%, respectively; **Fig. 8A**, *right*) and Tib nerve stimulation (F_(2,46)_ = 6.30, *p* = 0.004, −82%; F_(2,60)_ = 7.77, *p* = 0.001, −111%, respectively; **Fig. 8B**, *right*). Note that stimulating the Tib nerve before the perineal stimulation evoked P1 responses in 77% of the trials whereas during pinch of the perineal region we observed N1 responses in 77% of the trials, which explains why the modulation exceeded 100%.

**Figure 8.**
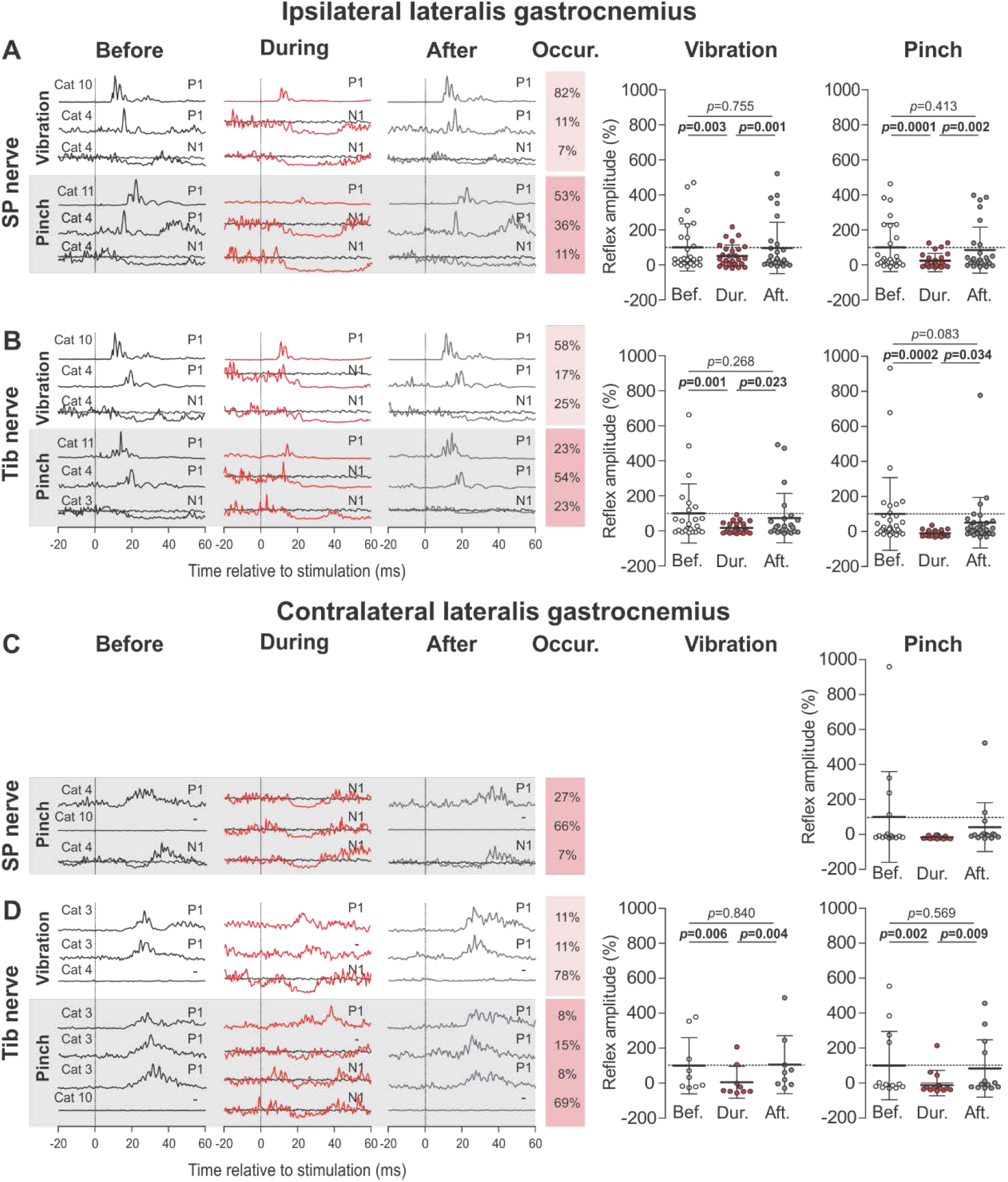
Modulation of reflex responses in the ipsilateral and contralateral lateral gastrocnemius muscle by mechanical stimulation of the perineal region. Short-latency ipsilateral and contralateral reflex responses evoked by stimulating the superficial peroneal (SP) (panels ***A*** and ***C***) and tibial (Tib) nerves (***B*** and ***D***, respectively) were showed. *Left*: All reflex responses patterns are illustrated for each condition. For each pattern, waveforms are averages of 13-15 stimulations per period for a representative cat in an 80-ms window. To better visualize N1 responses, we superimposed averaged traces that received a stimulation (line in black, red or grey) with averaged traces without stimulation (line in dark grey). On the right of each pattern, the occurrence (occur.) of the reflex responses pattern in percentage are illustrated. *Right*: the effect of vibration and pinch of the perineal region are illustrated for each period (Bef., before; Dur., during; Aft., after). Scatterplots show the mean ± SD of short-latency responses from pooled data (ipsilateral SP-vibration *n* = 28 trials from 6 cats; ipsilateral and contralateral SP-pinch *n* = 28 and 15 trials from 6 and 4 cats, respectively; ipsilateral and contralateral Tib-vibration *n* = 24 and 9 trials from 5 and 3 cats, respectively; ipsilateral and contralateral Tib-pinch *n* = 31 and 13 trials from 7 and 5 cats, respectively). *p* values comparing periods were obtained with Fisher’s post-hoc test if a significant main effect of the ANOVA was found.

In the contralateral LG, stimulating the SP nerve during vibration did not evoke enough reflex responses to perform statistical analysis. In the other conditions, perineal stimulation mainly caused the appearance of N1 responses with SP or Tib nerve stimulation when no (66-78% of trials) or P1 (8-27% of trials) responses were observed before and after. Vibration and pinch of the perineal region significantly decreased reflex responses evoked by Tib nerve stimulation (F_(2,16)_ = 6.92, *p* = 0.007, −94%; F_(2,24)_ = 6.73, *p* = 0.005, −100%, respectively; **Fig. 8D**, *right*) whereas pinch had no significant effect on responses evoked by stimulating the SP nerve (F_(2,28)_ = 2.84, *p* = 0.076; **Fig. 8C**, *right*).

In summary, mechanically stimulating the perineal region with vibration or pinch decreased ipsilateral and contralateral reflex responses evoked by SP or Tib nerve stimulation in all studied muscles, as summarized in **Table 3**. In some cases, perineal stimulation also caused the appearance of inhibitory responses or a switch from a positive to a negative response. Overall, pinch (*n* = 8) of the perineal region produced a greater number of significant decreases in ipsilateral reflex responses compared to vibration (*n* = 3).

**Table 3.** Summary of reflex modulations by vibration or pinch of the perineal region. For each muscle, ↑ or ↓ represent a significant increase or decrease, respectively, in reflex response during vibration/pinch of the perineal region. The percentage change is indicated in parentheses. ↔ represent a non-significant difference in reflex response with vibration/pinch of the perineal region. An empty box means an insufficient number of reflex responses to perform statistical analyses. SP, superficial peroneal nerve; Tib, tibial nerve.

|  | Vibration |  | Pinch |  |
| --- | --- | --- | --- | --- |
|  | SP | Tib | SP | Tib |
| I-St | ↔ | ↔ | ↓ (-30%) | ↓ (-33%) |
| I-IP | ↔ | ↔ | ↓ (-53%) | ↓ (-55%) |
| I-VL | ↔ | ↓ (-77%) | ↓ (-93%) | ↓ (-109%) |
| I-LG | ↓ (-48%) | ↓ (-82%) | ↓ (-75%) | ↓ (-111%) |
| C-St |  | ↓ (-129%) | ↓ (-193%) | ↓ (-131%) |
| C-IP |  | ↓ (-47%) |  | ↓ (-73%) |
| C-VL |  | ↓ (-70%) | ↓ (-98%) | ↓ (-87%) |
| C-LG |  | ↓ (-94%) | ↔ | ↓ (-100%) |

## Discussion

We investigated the effects of mechanically stimulating the perineal region on rhythmic activity and on short-latency cutaneous reflex responses in ipsilateral and contralateral hindlimb muscles of chronic spinal cats. Not surprisingly, perineal stimulation effectively facilitated or triggered unilateral and bilateral rhythmic hindlimb activity. Adding perineal stimulation to nerve stimulations increased the frequency and amplitude of rhythmic bursts in ipsilateral flexors and extensors and disrupted the timing of the rhythm in relation to nerve stimulation. However, contrary to our hypothesis, we observed that vibration and pinch of the perineal region decreased ipsilateral and contralateral reflex responses evoked by SP or Tib nerve stimulation in flexors and extensors. Interestingly, perineal stimulation could change the type of reflex response evoked by nerve stimulations, from positive to negative responses. Therefore, our results indicate that mechanoreceptive inputs from the perineal region do not mediate their excitatory effect on locomotion and weight support by increasing the gain of short-latency cutaneous reflexes from the foot. However, perineal stimulation does induce a state that modulates reflex response patterns.

### Perineal stimulation facilitates and modulates rhythmic activity initiated by nerve stimulation in hindlimb muscles

As recently shown (Merlet *et al*., 2020), we observed that stimulating the SP or Tib nerves generates rhythmic bursts of activity in some hindlimb muscles. Bursts in ipsilateral flexors were initiated with the nerve stimulation (phasing ~1) whereas bursts in ipsilateral extensors occurred spontaneously, most likely by engaging the spinal locomotor CPG, but ceased with the following nerve stimulation that initiated another flexor burst. In other words, electrical nerve stimulation reset the extension phase to flexion and entrained the rhythm. While the SP nerve at ankle level is entirely cutaneous, the Tib nerve is mixed, although its effect is generally attributed to cutaneous afferents (Loeb, 1993; Frigon & Rossignol, 2008). Resetting and entrainment of the rhythm indicate that cutaneous afferents of the SP and Tib nerves have direct access to the rhythm generating portion of the spinal locomotor CPG (reviewed in Hultborn *et al*., 1998; Pearson *et al*., 1998; Rossignol *et al*., 2006; McCrea & Rybak, 2008; Gossard *et al*., 2011). We propose that the rhythm induced by nerve stimulation corresponds to a weak locomotor-like state.

On the other hand, adding perineal stimulation to SP and Tib nerve stimulations modulated rhythmic activity by decreasing cycle and burst durations and increasing mean EMG amplitude of flexors and extensors. This corresponds to a strong locomotor-like state. Perineal stimulation also disrupted the timing of flexor burst onset to the nerve stimulation. Thus, without perineal stimulation, phasic sensory inputs from the SP and Tib nerves initiate and entrain a relatively weak rhythm. Tonic sensory feedback from the perineal stimulation generates a stronger rhythm and phasic sensory inputs have more difficulty entraining this new rhythm. With perfect entrainment, nerve stimulations at 0.5 Hz would have generated a rhythm with a cycle of 2 s. During treadmill locomotion, this corresponds to a speed of ~0.1 m/s in spinal cats (Frigon *et al*., 2017), which is a slow stepping speed and spinal cats do not step consistently at this speed (Dambreville *et al*., 2015). Intact cats cannot consistently perform treadmill locomotion below 0.3 m/s. The mean cycle duration that we observed before perineal stimulation was ~1.4-1.8 s and perineal stimulation reduced these values to ~1.0 s, corresponding to a treadmill speed of ~0.4-0.5 m/s, which is within the lower range of preferred stepping speeds on a treadmill in intact cats (Frigon, 2011). To recapitulate, in our preparation, the spinal locomotor CPG received two types of sensory inputs, phasic inputs from the nerve stimulations and a tonic input from the perineal stimulation. Perineal stimulation had a potent effect on the spinal locomotor CPG and brought its rhythmicity closer to its normal frequency, thus disrupting the entrainment from the phasic sensory inputs of the nerve stimulations.

The present study showed that vibration or pinch of the perineal region was equally effective in triggering unilateral and bilateral rhythmic activity (**Table 1**). Vibration can activate a variety of mechanoreceptors in muscle and skin, while the pinch activates only cutaneous mechanoreceptors. Thus, it is likely that the effect of perineal stimulation is primarily, if not exclusively, mediated by cutaneous inputs. What type of cutaneous afferents from the perineal region mediate the excitatory effect on the spinal locomotor CPG? Although we can only speculate, Viala *et al*. (1978) showed that low-threshold Aα, Aβ and C afferents from the lumbosacral skin had an excitatory effect on the spinal locomotor CPG of curarized decerebrate rabbits. In contrast, Aδ fibers were highly inhibitory and when mechanically stimulating the lumbosacral skin, the net effect was an inhibition of weight support and locomotion, as shown in cats and rabbits (Viala & Buser, 1974; Viala *et al*., 1978; Frigon *et al*., 2012; Hurteau *et al*., 2015; Merlet *et al*., 2020). It is possible that Aα, Aβ and C cutaneous afferents from the perineal region are also excitatory, while Aδ fibers are inhibitory, but when the population is activated the net excitatory effect is stronger than the inhibitory one. However, this hypothesis remains to be investigated. We also do not know if phasic sensory inputs to the pudendal nerve, which supplies the perineal skin, can entrain and reset the rhythm.

Interestingly, we also observed that perineal stimulation induced different patterns of rhythmic activity between the ipsilateral and contralateral sides relative to nerve stimulation. The presence of various relationships in the number of cycles between the left and right hindlimbs indicates that the fast side displayed one, two, three, four, five and even six cycles for every cycle of the slow side. While the 1:1 relationship predominated, we observed 20-40% of 1:2+ relationships. These types of relationships are reminiscent of split-belt locomotion, with the slow limb stepping at a slow speed, such as 0.1 or 0.2 m/s (Frigon et al. 2017). In our preparation, with the hindlimbs restrained, the slow and fast limbs could be either ipsilateral or contralateral to the nerve stimulation. Thus, it does not seem that nerve stimulation triggered these events. In some cases, we also observed that the duration and amplitude of the last extensor burst of the fast limb was increased. This phenomenon has been reported during split-belt locomotion and likely facilitates the transition from stance to swing for the slow limb in order to maintain stable locomotion when one hindlimb takes more steps than the other (Frigon *et al*., 2017). Thus, even though the hindlimbs were restrained, the pattern of EMG activity appeared functional to aid in the transfer of load from one limb to another. What triggers these types of relationships? Sensory feedback is important in regulating the timing of phase transitions and likely generates these relationships, in a sense ‘misleading’ the spinal locomotor CPG. It was shown that stretch-sensitive inputs from hip flexor and hip extensor muscles and load-sensitive inputs from extensor muscles and cutaneous mechanoreceptors of the plantar surface influence phase transitions in cats and humans (reviewed in Duysens *et al*., 2000; Rossignol *et al*., 2006; Pearson, 2008). Although the hindlimbs are restrained in our preparation, the rhythmic activity caused by perineal stimulation produce muscle contractions and group I/II afferents from Golgi tendon organs and muscle spindles are undoubtedly activated, thus influencing phase transitions. Whatever the mechanism, the spinal locomotor CPG seems to interpret the sensory feedback as consistent with one limb stepping faster than the other does.

### Perineal stimulation decreases the gain of cutaneous reflexes from the foot and induces reflex reversals

Our results showed that mechanically stimulating the perineal region with vibration or pinch decreased short-latency cutaneous reflex responses in ipsilateral and contralateral hindlimb muscles. The reduction in reflex gain during perineal stimulation was independent of background EMG activity preceding the sensory volley from the electrical nerve stimulation, consistent with premotoneuronal mechanisms. Premotoneuronal mechanisms can include gating of the transmission by interneurons within or outside the reflex pathway or the primary afferent itself. Presynaptic inhibition of SP or Tib nerve afferents is a likely mechanism for reduced reflex responses, because it was observed in all muscles ipsilateral and contralateral to the nerve stimulation during vibration and pinch of the perineal region. The simplest explanation for this generalized decrease is reduced neurotransmitter released from primary afferent terminals (Burke, 1999; Rudomin & Schmidt, 1999). One scenario is that the spinal locomotor CPG sets the level of presynaptic inhibition of primary afferents on a task-dependent basis. However, this does not explain the appearance of N1 responses or the switch from P1 to N1 responses. Stimulating the SP or Tib nerves during locomotion in cats and humans is known to induce excitatory and inhibitory responses, depending on the muscle and the phase (Forssberg & Grillner, 1973; Abraham *et al*., 1985; Yang & Stein, 1990; Duysens *et al*., 1990; Frigon & Rossignol, 2007; Frigon *et al*., 2009; Hurteau *et al*., 2018). Thus, SP and Tib afferents activate reflex pathways that end on both last-order excitatory and inhibitory interneurons. The expression of one pathway over the other is due to the balance of excitation and inhibition in these alternate pathways. In our study, the switch from P1 to N1 responses could have been mediated by the spinal locomotor CPG and/or by perineal inputs that change the balance in favor of the inhibitory pathway. **Figure 9** illustrates potential mechanism involved in the modulation of cutaneous reflexes in the LG muscle evoked by left SP nerve stimulation with perineal stimulation.

**Figure 9.**
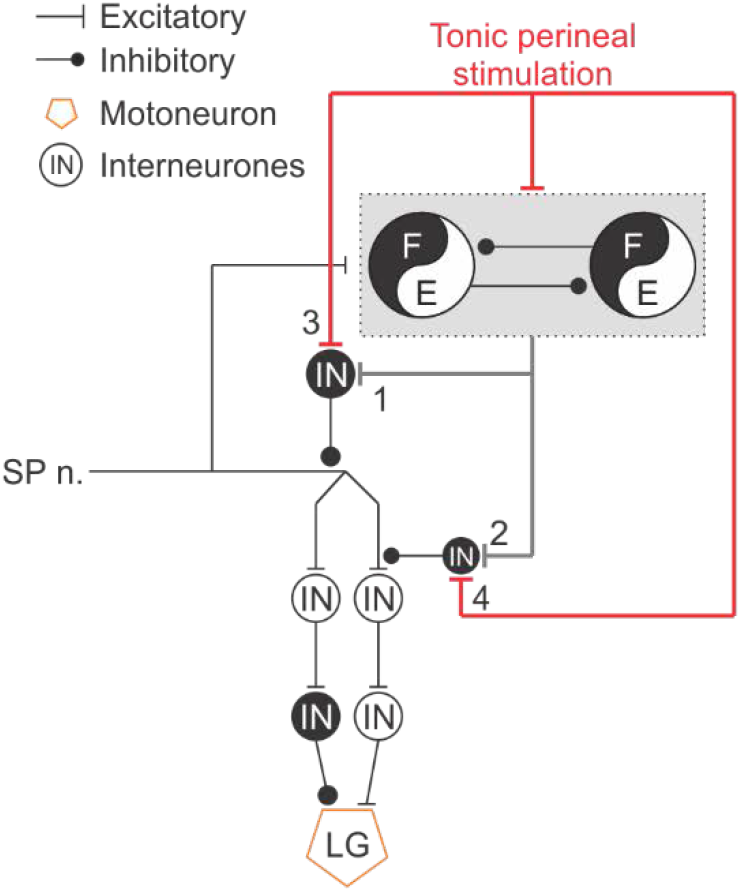
Schematic illustration of potential mechanisms modulating cutaneous reflexes with perineal stimulation. Some potential mechanisms involved in the modulation of cutaneous reflexes modulation in the lateralis gastrocnemius (LG) muscle evoked by left superficial peroneal nerve (SP.n) stimulation with perineal stimulation are illustrated. 1: The central pattern generator (CPG) sets the level of presynaptic inhibition of the primary afferent. 2: The CPG independently controls the excitability of different reflex pathways. 3: Perineal inputs directly regulate presynaptic inhibition of the primary afferent. 4: Perineal inputs independently control the excitability of different reflex pathways. E, extensor; F, flexor; IN, interneurons.

Task- and phase-dependent modulation of H-reflexes and cutaneous reflexes have been shown in several studies in humans (reviewed in Zehr & Stein, 1999). In cats, cutaneous reflexes are also modulated powerfully in a phase- and task-dependent manner during locomotion (Duysens & Loeb, 1980; Abraham *et al*., 1985; Frigon & Rossignol, 2007; Frigon *et al*., 2009; Hurteau *et al*., 2018, 2017; Hurteau & Frigon, 2018). For example, in humans, the H-reflex is reduced during walking compared to standing (Capaday & Stein, 1986) or during running compared to walking (Capaday & Stein, 1987) whereas cutaneous reflexes are greater during running than during standing (Duysens *et al*., 1993). Interestingly, studies in arm (Zehr & Kido, 2001) and leg (Zehr *et al*., 2001) muscles reported a switch in the sign of middle latency cutaneous reflexes, from facilitation during static contraction to inhibition during movement. Our results also highlighted a switch from P1 response during the “static state” to N1 response during the “locomotor state” (i.e. with perineal stimulation). Therefore, in our preparation, perineal stimulation induces a locomotor-like state that modulates the response pattern of cutaneous reflex.

The increase in spinal neuronal excitability provided by tonic perineal stimulation could replace monoaminergic drive from the brainstem (Harnie *et al*., 2019), which is known to facilitate hindlimb locomotion in spinal mammals (Forssberg & Grillner, 1973; Barbeau *et al*., 1987; Barbeau & Rossignol, 1990; Chau *et al*., 1998a, 1998b; Feraboli-Lohnherr *et al*., 1999; Kim *et al*., 2001; Antri *et al*., 2002, 2005; Sławińska *et al*., 2012). It was also reported that the injection of noradrenergic or serotonergic agonists differently influences the excitability of cutaneous reflexes as well as the modulation of a robustly recovered hindlimb locomotor pattern in late-spinal cats stepping on a treadmill (Barbeau *et al*., 1987; Barbeau & Rossignol, 1990, 1991; Chau *et al*., 1998b). Indeed, injection of an α-2 noradrenergic agonist, clonidine, produces a decrease in cutaneous reflexes excitability without apparent changes in the mean amplitude of flexor and extensor bursts (Barbeau *et al*., 1987; Chau *et al*., 1998b) whereas injection of a serotonergic agonist, 5-hydroxytryptamine (5-HT) increases cutaneous reflexes excitability and the mean amplitude of flexor and extensor (Barbeau & Rossignol, 1990). Both types of drug increased cycle duration (Barbeau *et al*., 1987; Barbeau & Rossignol, 1990, 1991; Chau *et al*., 1998b). When comparing our results and considering the role of the noradrenergic system in initiating and modulating hindlimb locomotion in spinal cats (Barbeau *et al*., 1987; Barbeau & Rossignol, 1991; Chau *et al*., 1998b) as well as the reduction of cutaneous reflexes (Barbeau *et al*., 1987; Chau *et al*., 1998b), we can speculate that tonic perineal stimulation acted through a similar mechanism, albeit with some differences, as we observed a reduction in cycle duration.

It is entirely possible and even likely that the reflex modulation that we observed in spinal cats is different from what we would observe in intact cats. After a complete spinal cord injury, cutaneous reflexes are modified because of changes in the balance between excitation and inhibition with spinal sensorimotor circuits (reviewed in Frigon & Rossignol, 2006; Rossignol & Frigon, 2011), and/or by expansion of cutaneous receptive fields (Schouenborg *et al*., 1992; Andersen *et al*., 2004). For example, stimulating cutaneous nerves during stance normally evokes N1 responses in ipsilateral extensors of intact cats (Duysens & Loeb, 1980; Abraham *et al*., 1985; Frigon & Rossignol, 2007, 2008; Frigon *et al*., 2009; Hurteau *et al*., 2018). However, after spinal transection, P1 responses to the same nerve stimulation, in the same animal, are frequently observed (Frigon & Rossignol, 2008).

Interestingly, we reported that pinching the skin of the perineal region was more effective than vibration in decreasing the gain of cutaneous reflexes (**Table 3**). As the afferents activated by vibration and pinch differ (Merlet *et al*., 2020), we propose that the greater effect of pinch in decreasing the gain of short-latency cutaneous reflexes was due to the specificity of afferents involved (i.e. solely cutaneous afferents).

### Concluding remarks

A better understanding of how sensory feedback from the perineal region interacts with spinal sensorimotor circuits is important because it has potent effects on weight support and locomotor activity in mammals with a complete spinal cord injury (Barbeau & Rossignol, 1987; Belanger *et al*., 1996; Leblond *et al*., 2003; Langlet *et al*., 2005; Hochman, 2012; Alluin *et al*., 2015; Harnie *et al*., 2019). Our results show that perineal stimulation induces a state that reorganizes or reduces reflex pathways. From a functional perspective, why would it be useful to reduce cutaneous reflexes during perineal stimulation? The functional purpose of the pathway from the perineal region to the spinal locomotor CPG is not clear but it could play an important survival function, such as increasing locomotor speed when a predator contacts this sensitive area (Rossignol *et al*., 2006). Jankowska (2008, 2015) argued that the role of the spinal circuitry is to increase or decrease specific aspects of somatosensory inputs depending on the demands of a given situation. In this context, the spinal locomotor CPG might reduce the gain of cutaneous reflexes to lower the effect of excitatory inputs in this more excitable locomotor state, as the animal has switched from an exploratory to an escape behavior.

## Author’s contributions

All authors had full access to all the data in the study and take responsibility for the integrity of the data and the accuracy of the data analysis. *Conceptualization*, J.H. and A.F.; *Methodology*, J.H. and A.F.; *Validation*, A.N.M., J.H., M.M. and A.F.; *Formal Analysis*, A.N.M. and M.M.; *Investigation*, J.H., A.D. and A.F;. *Resources*, N.G. and A.F.; *Writing – Original Draft Preparation*, A.N.M. and A.F.; *Writing – Review & Editing*, A.N.M., J.H., M.M., A.D., N.G. and A.F.; *Visualization*, A.N.M., J.H., and M.M.; *Supervision*, A.F.; *Funding Acquisition*, N.G. and A.F.

## Other Acknowledgments

We thank Philippe Drapeau (Rossignol and Drew laboratories, Université de Montréal) for providing data acquisition and analysis software.

## Data Accessibility links

The data that support the findings of this study are available from the corresponding author upon request.

## Literature Cited

Abraham LD, Marks WB & Loeb GE (1985). The distal hindlimb musculature of the cat: Cutaneous reflexes during locomotion. Exp Brain Res 58, 594–603.

Alluin O, Delivet-Mongrain H & Rossignol S (2015). Inducing hindlimb locomotor recovery in adult rat after complete thoracic spinal cord section using repeated treadmill training with perineal stimulation only. Journal of Neurophysiology 114, 1931–1946.

Andersen OK, Finnerup NB, Spaich EG, Jensen TS & Arendt-Nielsen L (2004). Expansion of nociceptive withdrawal reflex receptive fields in spinal cord injured humans. Clinical Neurophysiology 115, 2798–2810.

Antri M, Barthe J-Y, Mouffle C & Orsal D (2005). Long-lasting recovery of locomotor function in chronic spinal rat following chronic combined pharmacological stimulation of serotonergic receptors with 8-OHDPAT and quipazine. Neuroscience Letters 384, 162–167.

Antri M, Orsal D & Barthe J-Y (2002). Locomotor recovery in the chronic spinal rat: effects of long-term treatment with a 5-HT 2 agonist: 5-HT induced motor function recovery in spinal rat. European Journal of Neuroscience 16, 467–476.

Barbeau H, Julien C & Rossignol S (1987). The effects of clonidine and yohimbine on locomotion and cutaneous reflexes in the adult chronic spinal cat. Brain Research 437, 83–96.

Barbeau H & Rossignol S (1987). Recovery of locomotion after chronic spinalization in the adult cat. Brain Research 412, 84–95.

Barbeau H & Rossignol S (1990). The effects of serotonergic drugs on the locomotor pattern and on cutaneous reflexes of the adult chronic spinal cat. Brain Research 514, 55–67.

Barbeau H & Rossignol S (1991). Initiation and modulation of the locomotor pattern in the adult chronic spinal cat by noradrenergic, serotonergic and dopaminergic drugs. Brain Research 546, 250–260.

Belanger M, Drew T, Provencher J & Rossignol S (1996). A comparison of treadmill locomotion in adult cats before and after spinal transection. Journal of Neurophysiology 76, 471–491.

Bennett DJ, De Serres SJ & Stein RB (1996). Gain of the triceps surae stretch reflex in decerebrate and spinal cats during postural and locomotor activities. The Journal of Physiology 496, 837–850.

Bernard G, Bouyer L, Provencher J & Rossignol S (2007). Study of Cutaneous Reflex Compensation During Locomotion After Nerve Section in the Cat. Journal of Neurophysiology 97, 4173–4185.

Burke RE (1999). The use of state-dependent modulation of spinal reflexes as a tool to investigate the organization of spinal interneurons. Experimental Brain Research 128, 263–277.

Burke RE, Fedina L & Lundberg A (1971). Spatial synaptic distribution of recurrent and group Ia inhibitory systems in cat spinal motoneurones. The Journal of Physiology 214, 305–326.

Capaday C & Stein R (1986). Amplitude modulation of the soleus H-reflex in the human during walking and standing. J Neurosci 6, 1308–1313.

Capaday C & Stein RB (1987). Difference in the amplitude of the human soleus H reflex during walking and running. The Journal of Physiology 392, 513–522.

Cha J, Heng C, Reinkensmeyer DJ, Roy RR, Edgerton VR & De Leon RD (2007). Locomotor Ability in Spinal Rats Is Dependent on the Amount of Activity Imposed on the Hindlimbs during Treadmill Training. Journal of Neurotrauma 24, 1000–1012.

Chau C, Barbeau H & Rossignol S (1998a). Early Locomotor Training With Clonidine in Spinal Cats. Journal of Neurophysiology 79, 392–409.

Chau C, Barbeau H & Rossignol S (1998b). Effects of Intrathecal α 1 - and α 2-Noradrenergic Agonists and Norepinephrine on Locomotion in Chronic Spinal Cats. Journal of Neurophysiology 79, 2941–2963.

Crone C, Hultborn H, Jespersen B & Nielsen J (1987). Reciprocal Ia inhibition between ankle flexors and extensors in man. The Journal of Physiology 389, 163–185.

Dambreville C, Labarre A, Thibaudier Y, Hurteau M-F & Frigon A (2015). The spinal control of locomotion and step-to-step variability in left-right symmetry from slow to moderate speeds. Journal of Neurophysiology 114, 1119–1128.

De Leon RD, Hodgson JA, Roy RR & Edgerton VR (1999). Retention of Hindlimb Stepping Ability in Adult Spinal Cats After the Cessation of Step Training. Journal of Neurophysiology 81, 85–94.

Duysens J, Clarac F & Cruse H (2000). Load-Regulating Mechanisms in Gait and Posture: Comparative Aspects. Physiological Reviews 80, 83–133.

Duysens J & Loeb GE (1980). Modulation of ipsi- and contralateral reflex responses in unrestrained walking cats. Journal of Neurophysiology 44, 1024–1037.

Duysens J, Tax AAM, Trippel M & Dietz V (1993). Increased amplitude of cutaneous reflexes during human running as compared to standing. Brain Research 613, 230–238.

Duysens J, Trippel M, Horstmann GA & Dietz V (1990). Gating and reversal of reflexes in ankle muscles during human walking. Exp Brain Res 82, 351–8.

Forssberg H & Grillner S (1973). The locomotion of the acute spinal cat injected with clonidine i.v. Brain Research 50, 184–186.

Forssberg H, Grillner S, Halbertsma J & Rossignol S (1980). The locomotion of the low spinal cat. II. Interlimb coordination. Acta Physiologica Scandinavica 108, 283–295.

Frigon A (2011). Interindividual variability and its implications for locomotor adaptation following peripheral nerve and/or spinal cord injury. In Progress in Brain Research, pp. 101–118. Elsevier. Available at: https://linkinghub.elsevier.com/retrieve/pii/B9780444538253000127 [Accessed June 10, 2020].

Frigon A, Barrière G, Leblond H & Rossignol S (2009). Asymmetric Changes in Cutaneous Reflexes After a Partial Spinal Lesion and Retention Following Spinalization During Locomotion in the Cat. Journal of Neurophysiology 102, 2667–2680.

Frigon A, Desrochers É, Thibaudier Y, Hurteau M-F & Dambreville C (2017). Left-right coordination from simple to extreme conditions during split-belt locomotion in the chronic spinal adult cat: Left-right coordination during locomotion. J Physiol 595, 341–361.

Frigon A & Rossignol S (2006). Functional plasticity following spinal cord lesions. In Progress in Brain Research, pp. 231–398. Elsevier. Available at: https://linkinghub.elsevier.com/retrieve/pii/S0079612306570165 [Accessed June 10, 2020].

Frigon A & Rossignol S (2007). Plasticity of Reflexes From the Foot During Locomotion After Denervating Ankle Extensors in Intact Cats. Journal of Neurophysiology 98, 2122–2132.

Frigon A & Rossignol S (2008). Adaptive changes of the locomotor pattern and cutaneous reflexes during locomotion studied in the same cats before and after spinalization: Changes in reflexes during locomotion after spinalization. The Journal of Physiology 586, 2927–2945.

Frigon A, Thibaudier Y, Johnson MD, Heckman CJ & Hurteau M-F (2012). Cutaneous inputs from the back abolish locomotor-like activity and reduce spastic-like activity in the adult cat following complete spinal cord injury. Experimental Neurology 235, 588–598.

Gossard J-P, Sirois J, Noué P, Côté M-P, Ménard A, Leblond H & Frigon A (2011). The spinal generation of phases and cycle duration. In Progress in Brain Research, pp. 15–29. Elsevier. Available at: https://linkinghub.elsevier.com/retrieve/pii/B9780444538253000073 [Accessed June 10, 2020].

Harnie J, Doelman A, de Vette E, Audet J, Desrochers E, Gaudreault N & Frigon A (2019). The recovery of standing and locomotion after spinal cord injury does not require task-specific training. eLife 8, e50134.

Hochman S (2012). Enabling techniques for in vitro studies on mammalian spinal locomotor mechanisms. Front Biosci 17, 2158.

Hultborn H, Conway BA, Gossard J-P, Brownstone R, Fedirchuk B, Schomburg ED, Enriquez-Denton M & Perreault M-C (1998). How Do We Approach the Locomotor Network in the Mammalian Spinal Cord?a. Annals NY Acad Sci 860, 70–82.

Hurteau M-F & Frigon A (2018). A Spinal Mechanism Related to Left–Right Symmetry Reduces Cutaneous Reflex Modulation Independently of Speed During Split-Belt Locomotion. J Neurosci 38, 10314–10328.

Hurteau M-F, Thibaudier Y, Dambreville C, Chraibi A, Desrochers E, Telonio A & Frigon A (2017). Nonlinear Modulation of Cutaneous Reflexes with Increasing Speed of Locomotion in Spinal Cats. J Neurosci 37, 3896–3912.

Hurteau M-F, Thibaudier Y, Dambreville C, Danner SM, Rybak IA & Frigon A (2018). Intralimb and Interlimb Cutaneous Reflexes during Locomotion in the Intact Cat. J Neurosci 38, 4104–4122.

Hurteau M-F, Thibaudier Y, Dambreville C, Desaulniers C & Frigon A (2015). Effect of stimulating the lumbar skin caudal to a complete spinal cord injury on hindlimb locomotion. Journal of Neurophysiology 113, 669–676.

Jankowska E (1992). Interneuronal relay in spinal pathways from proprioceptors. Progress in Neurobiology 38, 335–378.

Kiehn O (2016). Decoding the organization of spinal circuits that control locomotion. Nat Rev Neurosci 17, 224–238.

Kim D, Murray M & Simansky KJ (2001). The Serotonergic 5-HT2C Agonist m-Chlorophenylpiperazine Increases Weight-Supported Locomotion without Development of Tolerance in Rats with Spinal Transections. Experimental Neurology 169, 496–500.

Langlet C, Leblond H & Rossignol S (2005). Mid-Lumbar Segments Are Needed for the Expression of Locomotion in Chronic Spinal Cats. Journal of Neurophysiology 93, 2474–2488.

Leblond H, L’Espérance M, Orsal D & Rossignol S (2003). Treadmill Locomotion in the Intact and Spinal Mouse. J Neurosci 23, 11411–11419.

de Leon RD, Hodgson JA, Roy RR & Edgerton VR (1998). Locomotor Capacity Attributable to Step Training Versus Spontaneous Recovery After Spinalization in Adult Cats. Journal of Neurophysiology 79, 1329–1340.

Loeb GE (1993). The distal hindlimb musculature of the cat: interanimal variability of locomotor activity and cutaneous reflexes. Exp Brain Res 96, 125–140.

Lovely RG, Gregor RJ, Roy RR & Edgerton VR (1986). Effects of training on the recovery of full-weight-bearing stepping in the adult spinal cat. Experimental Neurology 92, 421–435.

Lovely RG, Gregor RJ, Roy RR & Edgerton VR (1990). Weight-bearing hindlimb stepping in treadmill-exercised adult spinal cats. Brain Research 514, 206–218.

Lundberg A, Malmgren K & Schomburg ED (1977). Cutaneous facilitation of transmission in reflex pathways from Ib afferents to motoneurones. The Journal of Physiology 265, 763–780.

Matthews PB (1986). Observations on the automatic compensation of reflex gain on varying the pre-existing level of motor discharge in man. The Journal of Physiology 374, 73–90.

McCrea DA & Rybak IA (2008). Organization of mammalian locomotor rhythm and pattern generation. Brain Research Reviews 57, 134–146.

McCrea DA, Shefchyk SJ, Stephens MJ & Pearson KG (1995). Disynaptic group I excitation of synergist ankle extensor motoneurones during fictive locomotion in the cat. The Journal of Physiology 487, 527–539.

Merlet AN, Harnie J, Macovei M, Doelman A, Gaudreault N & Frigon A (2020). Mechanically stimulating the lumbar region inhibits locomotor-like activity and increases the gain of cutaneous reflexes from the paws in spinal cats. Journal of Neurophysiology 123, 1026–1041.

Pearson KG (2008). Role of sensory feedback in the control of stance duration in walking cats. Brain Research Reviews 57, 222–227.

Pearson KG, Jiang W & Ramirez JM (1992). The use of naloxone to facilitate the generation of the locomotor rhythm in spinal cats. Journal of Neuroscience Methods 42, 75–81.

Pearson KG, Misiaszek JE & Fouad K (1998). Enhancement and Resetting of Locomotor Activity by Muscle Afferentsa. Annals NY Acad Sci 860, 203–215.

Pearson KG & Rossignol S (1991). Fictive motor patterns in chronic spinal cats. Journal of Neurophysiology 66, 1874–1887.

Pierrot-Deseilligny E, Bergego C, Katz R & Morin C (1981). Cutaneous depression of Ib reflex pathways to motoneurones in man. Exp Brain Res 42, 351–61.

Rossignol S, Dubuc R & Gossard J-P (2006). Dynamic Sensorimotor Interactions in Locomotion. Physiological Reviews 86, 89–154.

Rossignol S & Frigon A (2011). Recovery of Locomotion After Spinal Cord Injury: Some Facts and Mechanisms. Annu Rev Neurosci 34, 413–440.

Rudomin P & Schmidt RF (1999). Presynaptic inhibition in the vertebrate spinal cord revisited. Experimental Brain Research 129, 1–37.

Schouenborg J, Holmberg H & Weng H-R (1992). Functional organization of the nociceptive withdrawal reflexes: II. Changes of excitability and receptive fields after spinalization in the rat. Exp Brain Res 90, 469–78.

Sherrington CS (1910). Flexion-reflex of the limb, crossed extension-reflex, and reflex stepping and standing. The Journal of Physiology 40, 28–121.

Shimamura M & Livingston RB (1963). Longitudinal conduction systems serving spinal and brain-stem coordination. Journal of Neurophysiology 26, 258–272.

Shurrager PS & Dykman RA (1951). Walking spinal carnivores. Journal of Comparative and Physiological Psychology 44, 252–262.

Sławińska U, Majczyński H, Dai Y & Jordan LM (2012). The upright posture improves plantar stepping and alters responses to serotonergic drugs in spinal rats: The upright posture improves plantar stepping. The Journal of Physiology 590, 1721–1736.

Viala G & Buser P (1974). [Inhibition of spinal locomotor activity by a special method of somatic stimulation in rabbits]. Exp Brain Res 21, 275–84.

Viala G, Orsal D & Buser P (1978). Cutaneous fiber groups involved in the inhibition of fictive locomotion in the rabbit. Exp Brain Res 33, 257–67.

Yang JF & Stein RB (1990). Phase-dependent reflex reversal in human leg muscles during walking. Journal of Neurophysiology 63, 1109–1117.

Zehr EP, Hesketh KL & Chua R (2001). Differential Regulation of Cutaneous and H-Reflexes During Leg Cycling in Humans. Journal of Neurophysiology 85, 1178–1184.

Zehr EP & Kido A (2001). Neural control of rhythmic, cyclical human arm movement: task dependency, nerve specificity and phase modulation of cutaneous reflexes. The Journal of Physiology 537, 1033–1045.

Zehr EP & Stein RB (1999). What functions do reflexes serve during human locomotion? Progress in Neurobiology 58, 185–205.

